# An efficient and robust laboratory workflow and tetrapod database for larger scale eDNA studies

**DOI:** 10.1101/345082

**Authors:** Jan Axtner, Alex Crampton-Platt, Lisa A. Hörig, Azlan Mohamed, Charles C.Y. Xu, Douglas W. Yu, Andreas Wilting

## Abstract

**Background:** The use of environmental DNA, ‘eDNA,’ for species detection via metabarcoding is growing rapidly. We present a co-designed lab workflow and bioinformatic pipeline to mitigate the two most important risks of eDNA: sample contamination and taxonomic mis-assignment. These risks arise from the need for PCR amplification to detect the trace amounts of DNA combined with the necessity of using short target regions due to DNA degradation.

**Findings:** Our high-throughput workflow minimises these risks via a four-step strategy: (1) technical replication with two PCR *replicates* and two *extraction replicates*; (2) using multi-markers (*12S, *16S*, CytB*); (3) a ‘twin-tagging,’ two-step PCR protocol;(4) use of the probabilistic taxonomic assignment method *PROTAX*, which can account for incomplete reference databases.

As annotation errors in the reference sequences can result in taxonomic mis-assignment, we supply a protocol for curating sequence datasets. For some taxonomic groups and some markers, curation resulted in over 50% of sequences being deleted from public reference databases, due to (1) limited overlap between our target amplicon and reference sequences; (2) mislabelling of reference sequences; (3) redundancy.

Finally, we provide a bioinformatic pipeline to process amplicons and conduct *PROTAX* assignment and tested it on an ‘invertebrate derived DNA’ (iDNA) dataset from 1532 leeches from Sabah, Malaysia. Twin-tagging allowed us to detect and exclude sequences with non-matching tags. The smallest DNA fragment (16S) amplified most frequently for all samples, but was less powerful for discriminating at species rank. Using a stringent and lax acceptance criteria we found 162 (stringent) and 190 (lax) vertebrate detections of 95 (stringent) and 109 (lax) leech samples.

**Conclusions:** Our metabarcoding workflow should help research groups increase the robustness of their results and therefore facilitate wider usage of e/iDNA, which is turning into a valuable source of ecological and conservation information on tetrapods.

## Introduction

Monitoring, or even detecting, elusive or cryptic species in the wild can be challenging. In recent years there has been a rise in the availability of cost-effective DNA-based methods made possible by advances in high-throughput DNA sequencing (HTS). One such method is eDNA metabarcoding, which seeks to identify the species present in a habitat from traces of ‘environmental DNA’ (eDNA) in substrates such as water, soil, or faeces. A variant of eDNA metabarcoding, known as ‘invertebrate-derived DNA’ (iDNA) metabarcoding, targets the genetic material of prey or host species extracted from copro-, sarco- or haematophagous invertebrates. Examples include tick [1] s, blow or carrion flies [2; 3; 4; 5], mosquitoes [6; 7; 8; 9] and leeches [10; 11; 12; 13]. Many of these parasites are ubiquitous, highly abundant, and easy to collect, making them an ideal source of biodiversity data, especially for terrestrial vertebrates that are otherwise difficult to detect [10; 14; 15]. In particular, the possibility for bulk collection and sequencing in order to screen large areas and minimise costs is attractive. However, most of the recent studies on iDNA studies focus on single-specimen DNA extracts and Sanger sequencing and thus are not making use of the advances of HTS and a metabarcoding framework for carrying out larger scale biodiversity surveys.

That said, e/iDNA metabarcoding also poses several challenges, due to the low quality and low amounts of target DNA available, relative to non-target DNA (including the high-quality DNA of the live-collected, invertebrate vector). In bulk iDNA samples comprised of many invertebrate specimens, this problem is further exacerbated by the variable time since each individual has fed, if at all, leading to differences in the relative amounts and degradation of target DNA per specimen. This makes e/iDNA studies similar to ancient DNA samples, which also pose the problem of low quality and low amounts of target DNA [16; 17]. The great disparity in the ratio of target to non-target DNA and the low overall amount of the former requires an enrichment step, which is achieved via the amplification of a short target sequence (amplicon) by polymerase chain reaction (PCR) to obtain enough target material for sequencing. However, this enrichment step can result in false positive species detections, either through sample cross-contamination or through volatile short PCR amplicons in the laboratory, and in false-negative results, through primer bias and low concentrations of template DNA. Although laboratory standards to prevent and control for such false results are well established in the field of ancient DNA, there are still no best-practice guidelines for e/iDNA studies, and thus few studies sufficiently account for such problems [18].

The problem is exacerbated by the use of ‘universal’ primers used for the PCR, which maximise the taxonomic diversity of the amplified sequences. This makes the method a powerful biodiversity assessment tool, even where little is known a *priori* about which species might be found. However, using such primers, in combination with low quality and quantity of target DNA, which often requires a high number of PCR cycles to generate enough amplicon products for sequencing, makes metabarcoding studies particularly vulnerable to false results [13; 19; 20]. The high number of PCR cycles, combined with the high sequencing depth of HTS, also increase the likelihood that contaminants are amplified and detected, possibly to the same or greater extent as some true-positive trace DNA. As e/iDNA have been proposed as tools to detect very rare and priority conservation species such as the Saola, *Pseudoryx nghetinhensis* [10], false detection might result in misdirected conservation activities worth several hundreds of thousands of US dollars like for the ivory-billed woodpecker where most likely false evidence of the bird’s existence have been overemphasized to shore up political and financial support for saving it [21]. Therefore, similar to ancient DNA studies, great care must be taken to minimise the possibility for cross-contamination in the laboratory and to maximise the correct detection of species through proper experimental and analytical design. Replication in particular is an important tool for reducing the incidence of false negatives and detection of false positives but the trade-off is increased cost, workload, and analytical complexity [19].

An important source of false positive species detections is the incorrect assignment of taxonomies to the millions of short HTS reads generated by metabarcoding. Although there has been a proliferation of tools focused on this step, most can be categorised into just three groups depending on whether the algorithm utilises sequence similarity searches, sequence composition models, or phylogenetic methods [22; 23; 24]. The one commonality among all methods is the need for a reliable reference database of correctly identified sequences, yet there are few curated databases currently appropriate for use in e/iDNA metabarcoding. Two exceptions are SILVA [25] for the nuclear markers SSU and LSU rRNA used in microbial ecology, and BOLD (Barcode of Life Database; citation) for the COI ‘DNA barcode’ region. For other loci, a non-curated database downloaded from the INSDC (International Nucleotide Sequence Database Collaboration, e.g. GenBank) is generally used. However, the INSDC places the burden for metadata accuracy, including taxonomy, on the sequence submitters, with no restriction on sequence quality or veracity. For instance, specimen identification is often carried out by non-specialists, which increases error rates, and common laboratory contaminant species (e.g. human DNA sequences) are sometimes submitted in lieu of the sample itself. The rate of sequence mislabelling in fungi has been assessed for GenBank where it was up to 20% [26] and it is an issue that is often neglected [27; 28]. For several curated microbial databases (Greengenes, LTP, RDP, SILVA), mislabelling rates have been estimated at between 0.2% and 2.5% [29]. Given the lack of professional curation it is likely that the true proportion of mislabelled samples in GenBank is somewhere between these numbers. Moreover, correctly identifying such errors is labour-intensive, so most metabarcoding studies simply base their taxonomic assignments on sequence-similarity searches of the whole INSDC database (e.g. with BLAST) [3; 10; 12] and thus can only detect errors if assignments are ecologically unlikely. Furthermore, reference sequences for the species that are likely to be sampled in e/iDNA studies are often underrepresented in or absent from these databases, which increases the possibility of incorrect assignment. For instance, fewer than 50% of species occurring in a tropical megadiverse rainforest are represented in Genbank (see findings below). When species-level matches are ambiguous, it might still be possible to assign a sequence to a higher taxonomic rank by using an appropriate algorithm such as Metagenome Analyzer’s (MEGAN) Lowest Common Ancestor [30] or *PROTAX* [31].

We present here a complete laboratory workflow and complementary bioinformatics pipeline, starting from DNA extraction to taxonomic assignment of HTS reads using a curated reference database. The laboratory workflow allows for efficient screening of hundreds of e/iDNA samples. The workflow includes (1) two *extraction replicates* are separated during DNA extraction, and each is sequenced in two *PCR replicates* (Fig. 1); (2) robustness of taxonomic assignment is improved by using up to three mitochondrial markers; (3) a ‘twin-tagged’, two-step PCR protocol prevents cross-sample contamination as no unlabelled PCR products are produced (Fig. 2) while also allowing for hundreds of PCR products to be pooled before costly Illumina library preparation; (4) our bioinformatics pipeline includes a standardized, automated, and replicable protocol to create a curated database, which allows updating as new reference sequences become available, and to be expanded to other amplicons. We provide scripts for processing raw sequence data to quality-controlled dereplicated reads and for taxonomic assignment of these reads using *PROTAX* [31], a probabilistic method that has been shown to be robust even when reference databases are incomplete [23; 4] (all scripts are available from URL https://github.com/alexcrampton-platt/screenforbio-mbc).

**Figure 1:**
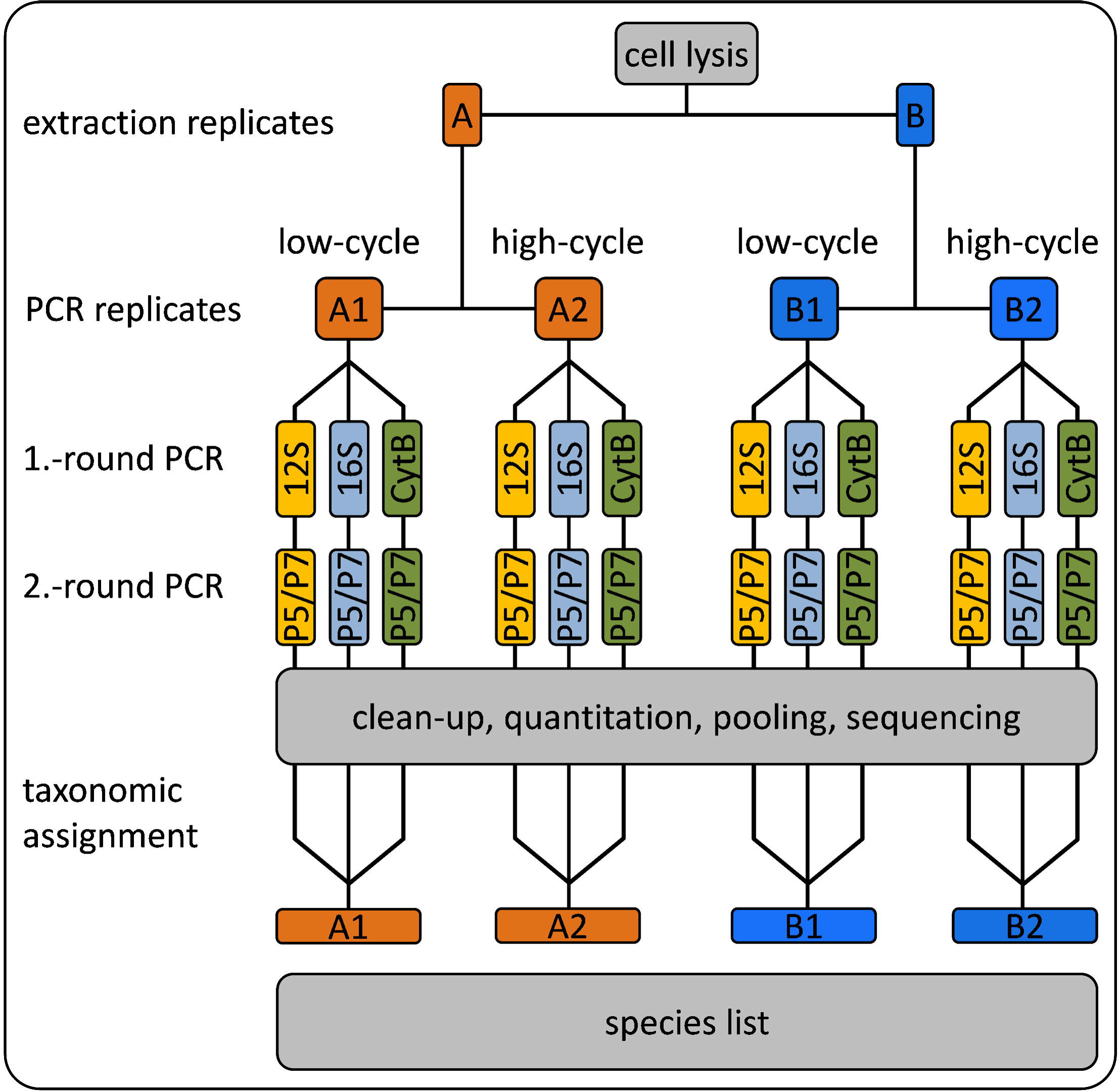
laboratory scheme; during DNA extraction the sample is split into two extraction replicates A & B. Our Protocol consists of two rounds of PCR that were the sample tags, the necessary sequencing primer and sequencing adapters are added to the the amplicons. For each extraction replicate we ran a low cycle PCR and a high cycle PCR for each marker that we have twelve independent *PCR replicates* per sample. All PCR products were sequenced and the obtained reads were taxonomically identified with *PROTAX*.

**Figure 2:**
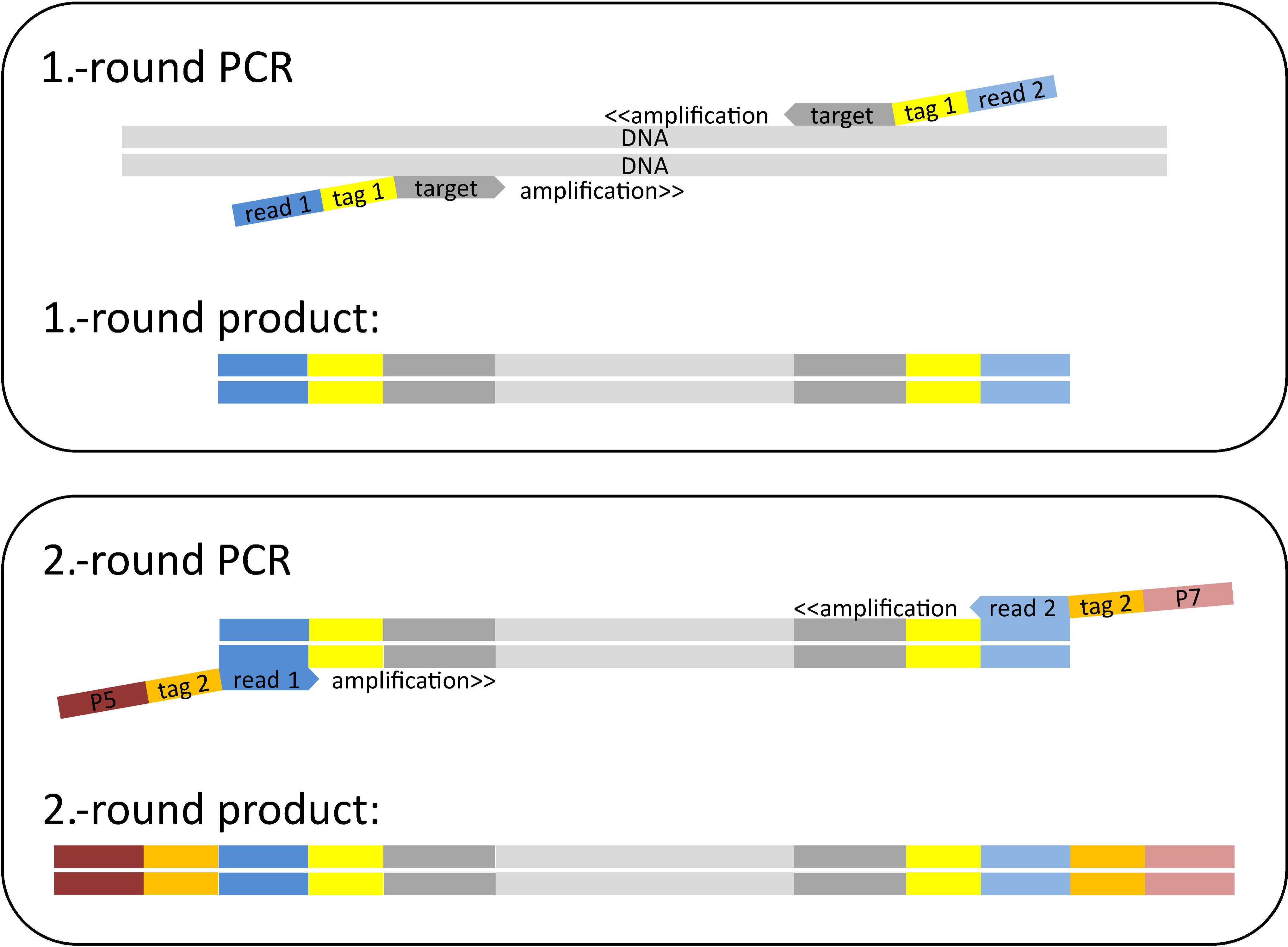
Scheme to build double ‘twin-tagged’ PCR libraries. The first round of PCR uses target-specific primers (*12S*, *16S*, or *CytB*, dark grey) that have both been extended with the same (i.e. ‘twin’) sample-identifying *tag* sequences *tag 1* (yellow) and then with the different *read 1* (dark blue) and *read 2* (light blue) sequence primers. The second round of PCR uses the priming sites of the *read 1* and *read 2* sequencing primers to add twin plate-identifying tag sequences *tag 2* (orange) and the P5 (dark red) and P7 (light red) Illumina adapters.

## Methods

### Establishment of the tetrapod reference database

#### Reference database

A custom bash script was written to generate a tetrapod reference database for up to four mitochondrial markers – a short 93 bp fragment of *16S* rRNA (*16S*), a 389 bp fragment of *12S* rRNA (*12S*), a 302 bp fragment of cytochrome b (*CytB*), and a 250 bp mitochondrial cytochrome c oxidase subunit I amplicon (*COI*) that has previously been used in iDNA studies [2]. An important time-saving step was the use of the FASTA-formatted Midori mitochondrial database [32], which is a lightly curated subset of Genbank. Our script updated the FASTA files with a subset of target species, removed errors and redundancy, trimmed the sequences to include only the amplicon regions, and output FASTA files with species names and GenBank accessions in the headers.

The script accepts four data inputs, two of which are optional. The required inputs are: (i) the Midori sequences (December 2015 ‘UNIQUE’, downloaded from http://www.reference-midori.info/download.php#) for the relevant genes and (ii) an initial reference taxonomy of tetrapods. This taxonomy is needed to find or generate a full taxonomic classification for each sequence because the taxonomies in Midori are from Genbank and thus include incorrect, synonymized, or incomplete taxonomies. Here we used the Integrated Taxonomic Information System (ITIS) classification for Tetrapoda, obtained with the R package *taxize* version 0.9.0 ([33], functions *downstream* and *classification*). The optional inputs are: (iii) supplementary FASTA files of reference sequences that should be added to the database, and (iv) a list of target species to be queried on GenBank to capture any sequences published since the December 2015 Midori dataset was generated.

For this study, 72 recently published [34) and 7 unpublished partial mitochondrial mammal genomes (Accession Numbers MH464789, MH464790, MH464791, MH464792, MH464793, MH464794, MH464795, MH464796, MH464797, MH464798, MH464799, MH464800, MH464801) were added as input (iii). A list of 103 mammal species known to be present in the sampling area plus *Homo sapiens* and our positive control *Myodes glareolus* was added as input (iv).

With the above inputs, the seven curation steps are: 1) remove sequences not identified to species; 2) add extra sequences from optional inputs (iii) and (iv) above; 3) trim the sequences to leave only the target amplicon; 4) remove sequences with ambiguities; 5) compare species names from the Midori dataset to the reference taxonomy from input (ii) and replace with a consensus taxonomy; 6) identify and remove putatively mislabeled sequences; 7) dereplicate sequences, retaining one haplotype per species.

The script is split into four modules, allowing optional manual curation at three key steps. The steps covered by each of the four modules are summarized in Table 2. The main programs used are highlighted and cited in the text where relevant, but many intermediate steps used common UNIX tools and unpublished lightweight utilities freely available from GitHub (Table 3).

##### Module 1

The first step is to select the tetrapod sequences from the Midori database for each of the four selected loci (input (i) above). This, and the subsequent step to discard sequences without strict binomial species names and reduce subspecies identifications to species-level, are made possible by the inclusion of the full NCBI taxonomic classification of each sequence in the FASTA header by the Midori pipeline. The headers of the retained sequences are then reformatted to include just the species name and GenBank accession separated by underscores. If desired, additional sequences from local FASTA files are now added to the Midori set (input (iii)). The headers of these FASTA files are required to be in the same format. Next, optional queries are made to the NCBI GenBank and RefSeq databases for each species in a provided list (input (iv)) for each of the four target loci, using NCBI’s Entrez Direct [35]. Matching sequences are downloaded in FASTA format, sequences prefixed as “UNVERIFIED” are discarded, the headers are simplified as previously, and those sequences not already in the Midori set are added. Trimming each sequence down to the relevant target marker was carried out in a two-step process in which *usearch* (*-search_pcr*) was used to select sequences where both primers were present, and these were in turn used as a reference dataset for *blastn* to select partially matching sequences from the rest of the dataset [36; 37]. Sequences with a hit length of at least 90% of the expected marker length were retained by extracting the relevant subsequence based on the BLAST hit co-ordinates. Sequences with ambiguous bases were discarded at this stage. In the final step in module 1, a multiple-sequence alignment was generated with MAFFT [38; 39] for each partially curated amplicon dataset (for the SATIVA step below). The script then breaks to allow the user to check for any obviously problematic sequences that should be discarded before continuing.

##### Module 2

The species labels of the edited alignments are compared with the reference taxonomy (input (ii)). Any species not found is queried against the Catalogue of Life database (CoL) via *taxize* in case these are known synonyms, and the correct species label and classification is added to the reference taxonomy. The original species label is retained as a key to facilitate sequence renaming, and a note is added to indicate its status as a synonym. Finally, the genus name of any species not found in the CoL is searched against the consensus taxonomy, and if found, the novel species is added by taking the higher classification levels from one of the other species in the genus. Orphan species labels are printed to a text file, and the script breaks to allow the user to check this list and manually create classifications for some or all if appropriate.

##### Module 3

This module begins by checking for any manually generated classification files (from the end of Module 2) and merging them with the reference taxonomy from Module 2. Any remaining sequences with unverifiable classifications are removed at this step. The next steps convert the sequences and taxonomy file to the correct formats for SATIVA [29], which detects possibly mislabelled sequences by generating a maximum likelihood phylogeny from the alignment in Module 1 and comparing each sequence’s taxonomy against its phylogenetic neighbors. Sequence headers in the edited MAFFT alignments are reformatted to include only the GenBank accession, and a taxonomy key file is generated with the correct classification listed for each accession number. In cases where the original species label is found to be a synonym, the corrected label is used. Putatively mislabeled sequences in each amplicon are then detected with SATIVA, and the script breaks to allow inspection of the results. The user may choose to make appropriate edits to the taxonomy key file or list of putative mislabels at this point.

##### Module 4

Any sequences that are still flagged as mislabelled at the start of the fourth module are deleted from the SATIVA input alignments, and all remaining sequences are relabelled with the correct species name and accession. A final consensus taxonomy file is generated in the format required by *PROTAX*. Alignments are subsequently unaligned prior to species-by-species selection of a single representative per unique haplotype. Sequences that are the only representative of a species are automatically added to the final database. Otherwise, all sequences for each species are extracted in turn, aligned with MAFFT, and collapsed to unique haplotypes with *collapsetypes_4.6.pl* (zero differences allowed; [40]). Representative sequences are then unaligned and added to the final database.

### iDNA samples

We used 242 collections of haematophagous terrestrial leeches from Deramakot Forest Reserve in Sabah, Malaysian Borneo stored in RNA fixating saturated ammonium sulfate solution as samples. Each sample consisted of one to 77 leech specimens (median 4). In total, 1532 leeches were collected, exported under the permit (JKM/MBS.1000-2/3 JLD.2 (8) issued by the Sabah Biodiversity Council), and analysed at the laboratories of the Leibniz-IZW.

### Laboratory workflow

The laboratory workflow is designed to both minimize the risk of sample cross-contamination and to aid identification of any instances that do occur. All laboratory steps (extraction, pre and post PCR steps, sequencing) took place in separate laboratories and no samples or materials were allowed to re-enter upstream laboratories at any point in the workflow. All sample handling was carried out under specific hoods that were wiped with bleach, sterilized, and UV irradiated for 30 minutes after each use. All labs are further UV irradiated for four hours each night.

#### DNA extraction

DNA was extracted from each sample in bulk. Leeches were cut into small pieces with a fresh scalpel blade and incubated in lysate buffer (proteinase K and ATL buffer at a ratio of 1:10; 0.2 ml per leech) overnight at 55 °C (12 hours minimum) in an appropriately sized vessel for the number of leeches (2 or 5 ml reaction tube). For samples with more than 35 leeches, the reaction volume was split in two and recombined after lysis.

Each lysate was split into two *extraction replicates* (A and B; maximum volume 600 μl) and all further steps were applied to these independently. We followed the DNeasy 96 Blood & Tissue protocol for animal tissues (Qiagen, Hilden -Germany) on 96 plates for cleanup. DNA was eluted twice with 100 μl TE buffer. DNA concentration was measured with PicoGreen dsDNA Assay Kit (Quant-iT, ThermoFisherScientific, Waltham -USA) in 384-well plate format using an appropriate plate reader (200 PRO NanoQuant, Tecan Trading AG, Männedorf - Switzerland). Finally, all samples were diluted to a maximum concentration of 10 ng/μl.

#### Two-round PCR protocol

We amplified three mitochondrial markers – a short 93 bp fragment of *16S* rRNA (*16S*), a 389 bp fragment of *12S* rRNA (*12S*), and a 302 bp fragment of cytochrome b (*CytB*). For each marker, we ran a two-round PCR protocol (Figs. 1, 2). The primers were chosen on the expectation of successful DNA amplification over a large number of tetrapod species [41; 42], and we tested the fit of candidate primers on an alignment of available mitochondrial sequences of 134 Southeast-Asian mammal species. Primer sequences are in Table 1.

**Table 1:**
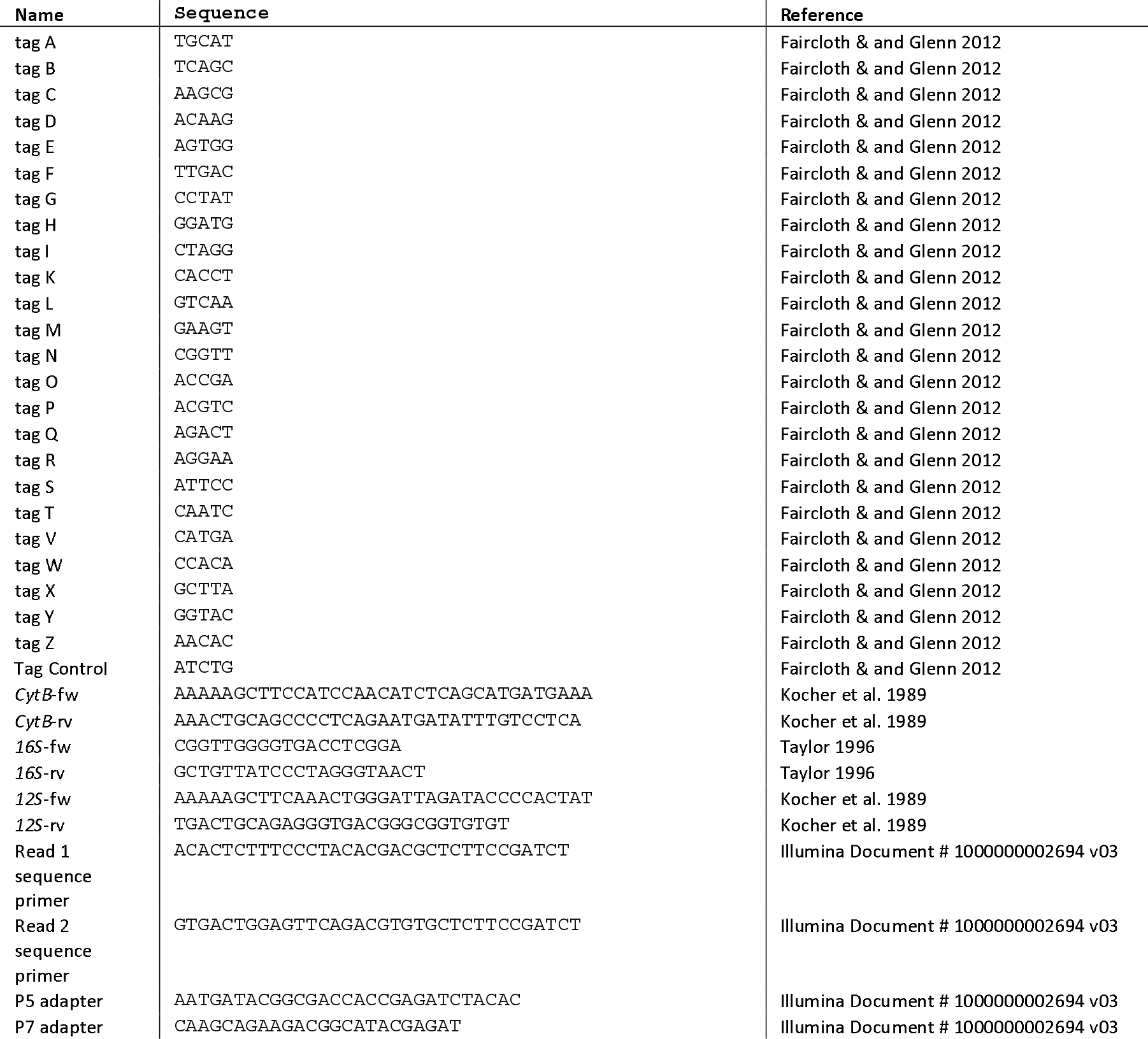
Sequence motifs that compose the 25 different target primers for the first and the second PCR. First PCR primers consist of target specific primer followed by an overhang out of sample specific *tag 1* and *read 1* and *read 2* sequencing primer, respectively. The second PCR primers consist of the read 1 or the read 2 sequencing primer followed by an plate specific tag 2 and the P5 and P7 adapters, respectively (see also Fig. 2).

**Table 2:**
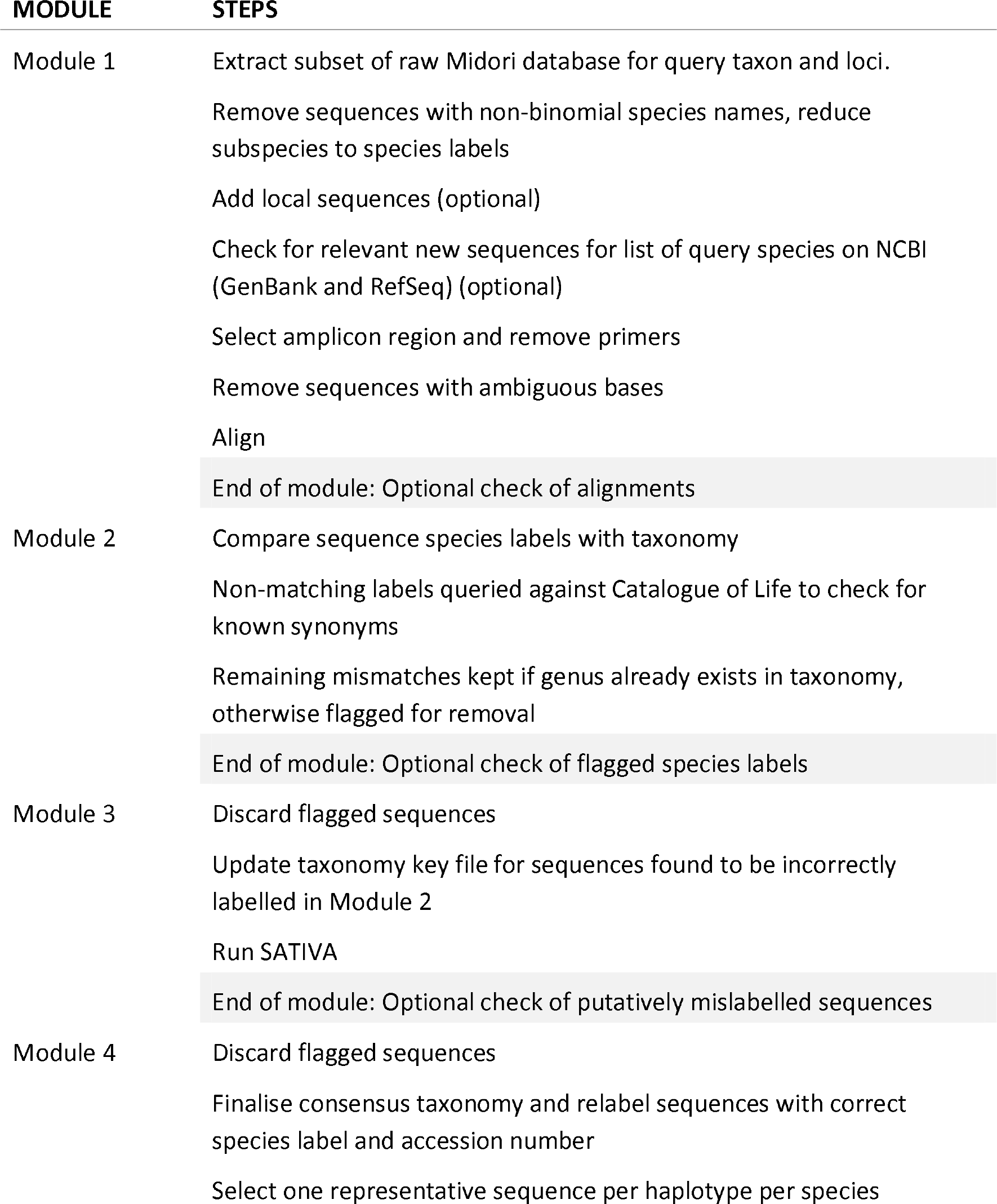
Main steps undertaken by each module of the database curation script.

**Table 3:** GNU core utilities and other lightweight tools used for manipulation of text and sequence files

| TOOL | FUNCTION | SOURCE |
| --- | --- | --- |
| awk, cut, grep,<br>join, sed, sort,<br>tr | Processing text files | GNU core utilities |
| seqbuddy | Processing FASTA/Q files | https://github.com/biologyguy/BuddySuite |
| seqkit | Processing FASTA/Q files | https://github.com/shenwei356/seqkit |
| seqtk | Processing FASTA/Q files | https://github.com/lh3/seqtk |
| tabtk | Processing tab-delimited<br>text files | https://github.com/lh3/tabtk |

##### Primer modification

We modified primers of the three markers to avoid the production of unlabelled PCR products, to allow the detection and deletion of tag-jumping events [43], and to reduce the cost of primers and library preparation. We used two rounds of PCR. The first round amplified the target gene and attached one of 25 different ‘twin-tag’ pairs (*tag 1*), identifying the sample within a given PCR. By ‘twin-tag,’ we mean that both the forward and reverse primers were given the *same* sample-identifying sequence (‘tags’) added as primer extensions (Fig. 2). The tags differed with a minimum pairwise distance of three nucleotides ([43]; Supplemental Table 1). These primers also contained different forward and reverse sequences (*Read 1 & Read 2 sequence primers*) (Supplemental Table 1) to act priming sites for the second PCR round (Fig. 2).

The second round added the Illumina adapters for sequencing and attached one of 20 twin-tag pairs (*tag 2*) identifying the PCR, with a minimum pairwise distance of three [44]. These primers also contained the Illumina P5 and P7 adapter sequences (Fig. 2). Thus no unlabelled PCR products were ever produced, and the combination of *tags 1* and 2 allowed the pooling of up to 480 (=24 × 20) samples in a single library preparation step (*one tag 1* was reserved for controls). Twin tags allowed us later to detect and delete tag jumping events [43] (Fig. 2).

##### Cycle number considerations

Because we know that our target DNA is at low concentration in the samples, we are faced with a trade-off between (1) using fewer PCR cycles (e.g. 30) to minimise amplification bias (caused by some target DNA binding better to the primer sequences and thus outcompeting other target sequences that bind less well [45]) and (2) using more PCR cycles (e.g. 40) to ensure that low-concentration target DNA is sufficiently amplified in the first place. Rather than choose between these two extremes, we ran both low- and high-cycle protocols and sequenced both sets of amplicons.

Thus, each of the two *extraction replicates* A and B was split and amplified using different cycle numbers (*PCR replicates* 1 and 2) for a total of four (*= 2 extraction replicates x 2 *PCR replicates* -> A1/A2 and B1/B2*) replicates per sample per marker (Fig. 1). For *PCR replicates* A1/B1, we used 30 cycles in the first PCR round to minimize the effect of amplification bias. For *PCR replicates* A2/B2, we used 40 cycles in the first PCR round to increase the likelihood of detecting species with very low input DNA (Fig. 1).

##### PCR protocol

The first-round PCR reaction volume was 20 μl, including 0.1 μM primer mix, 0.2 mM dNTPs, 1.5 mM MgCl2, 1x PCR buffer, 0.5 U AmpliTaq Gold™ (Invitrogen, Karlsruhe - Germany), and 2 μl of template DNA. Initial denaturation was 5 minutes at 95°C, followed by repeated cycles of 30 seconds at 95°C, 30 seconds at 54°C, and 45 seconds at 72°C. Final elongation was 5 minutes at 72°C. Samples were amplified in batches of 24 plus a negative (water) and a positive control (bank vole, *Myodes glareolus* DNA). All three markers were amplified simultaneously in individual wells for each batch of samples in a single PCR plate. Non-target by-products were removed as required from some *12S* PCRs by purification with magnetic Agencourt AMPure beads (Beckman Coulter, Krefeld -Germany).

In the second-round PCR, we used the same PCR protocol as above with 2 μl of the product of the first-round PCR and 10 PCR cycles.

#### Quality control and sequencing

Amplification was visually verified after the second-round PCR by gel electrophoresis on 1.5% agarose gels. Controls were additionally checked with a TapeStation 2200 (D1000 ScreenTape assay, Agilent, Waldbronn -Germany). All samples were purified with AMPure beads, using a bead-to-template ratio of 0.7:1 for *12S* and *CytB* products, and a ratio of 1:1 for *16S* products were always combined in a separate library. 12S/CytB libraries were sequenced independently from 16S libraries. Apart from our negative controls, we did not include samples that did not amplify, as this would have resulted in highly diluted libraries. Up to 11 libraries were sequenced on each run of Illumina MiSeq, following standard protocols. Libraries were sequenced with MiSeq Reagent Kit V3 (600 cycles, 300 bp paired-end reads) and had a final concentration of 11 pM spiked with 20 to 30% of PhiX control.

### Bioinformatics workflow

#### Read processing

Although the curation of the reference databases is our main focus, it is just one part of the bioinformatics workflow for e/iDNA metabarcoding. A custom bash script was used to process raw basecall files into demultiplexed, cleaned, and dereplicated reads in FASTQ format on a run-by-run basis. All runs and amplicons were processed with the same settings unless otherwise indicated. *bcl2fastq* (Illumina) was used to convert the basecall file from each library to paired-end FASTQ files, demultiplexed into the separate PCRs via the *tag 2* pairs, allowing up to 1 mismatch in each *tag 2*. Each FASTQ file was further demultiplexed into samples via the *tag 1* pairs using *AdapterRemoval* [46], again allowing up to 1 mismatch in each tag. These steps allowed reads to be assigned to the correct samples.

In all cases, amplicons were short enough to expect paired reads to overlap. For libraries with more than 1000 reads pairs were merged with *usearch* (*-fastq_mergepairs;* [47; 48]), and only successfully merged pairs were retained. For libraries with more than 500 merged pairs the primer sequences were trimmed away with *cutadapt* [49], and only successfully trimmed reads at least 90% of expected amplicon length were passed to a quality filtering step with *usearch* (*-fastq_filter*). Lastly, reads were dereplicated with usearch (*-derep_fulllength*), and singletons were discarded. The number of reads processed at each step for each sample are reported in a standard tab delimited txt-file.

#### Taxonomic assignment

The curated reference sequences and associated taxonomy were used for *PROTAX* taxonomic assignment of the dereplicated reads [24; 31]. *PROTAX* gives unbiased estimates of placement probability for each read at each taxonomic rank, allowing assignments to be made to a higher rank even when there is uncertainty at the species level. In other words, and unlike other taxonomic assignment methods, *PROTAX* can estimate the probability that a sequence belongs to a taxon that is not present in the reference database. This was considered an important feature due to the known incompleteness of the reference databases for tetrapods in the sampled location. As other studies have compared *PROTAX* with more established methods, e.g. MEGAN [30] (see [4; 24]), it was beyond the scope of this study to evaluate the performance of *PROTAX*.

Classification with *PROTAX* is a two-step process. Firstly, *PROTAX* selected a subset of the reference database that was used as training data to parameterise a *PROTAX* model for each marker, and secondly, the fitted models were used to assign four taxonomic ranks (species, genus, family, order) to each of the dereplicated reads, along with a probability estimate at each level. We also included the best similarity score of the assigned species or genus, mined from the LAST results (see below) for each read. This was helpful for flagging problematic assignments for downstream manual inspection, i.e. high probability assignments based on low similarity scores (implying that there are no better matches available) and low probability assignments based on high similarity scores (indicates conflicting database signal from several species with highly similar sequences).

Fitting the *PROTAX* model followed Somervuo et al. [31] except that 5000 training sequences were randomly selected for each target marker due to the large size of the reference database. In each case, 4500 training sequences represented a mix of known species with reference sequences (conspecific sequences retained in the database) and known species without reference sequences (conspecific sequences omitted, simulating species missing from the database), and 500 sequences represented previously unknown lineages distributed evenly across the four taxonomic levels (i.e. mimicked a mix of completely novel species, genera, families and orders). Pairwise sequence similarities of queries and references were calculated with LAST [50] following the approach of Somervuo et al. [31]. The models were weighted towards the Bornean mammals expected in the sampled area by assigning a prior probability of 90% to these 103 species and a 10% probability to all others ([31]; Supplemental Table 2). In cases of missing interspecific variation, this helped to avoid assignments to geographically impossible taxa, especially in case of the very short 93 bp fragment of *16S*. Maximum *a posteriori* (MAP) parameter estimates were obtained following the approach of Somervuo et al. [24], but the models were parameterised for each of the four taxonomic levels independently, with a total of five parameters at each level (four regression coefficients and the probability of mislabelling).

Dereplicated reads for each sample were then classified using a custom bash script on a run-by-run basis. For each sample, reads in FASTQ format were converted to FASTA, and pairwise similarities were calculated against the full reference sequence database for the applicable marker with LAST. Assignments of each read to a taxonomic node based on these sequence similarities were made using a Perl script and the trained model for that level. The taxonomy of each node assignment was added with a second Perl script for a final table including the node assignment, probability, taxonomic level, and taxonomic path for each read. Read count information was included directly in the classification output via the size annotation added to the read headers during dereplication. All Perl scripts to convert input files into the formats expected by *PROTAX*, R code for training the model following Somervuo et al. [31], and Perl scripts for taxonomic assignment were provided by P. Somervuo (personal communication).

#### Acceptance criteria

In total we had twelve PCR reactions per sample: two *extraction replicates A and B* X two *PCR replicates* 1 and 2 per extraction replication X the three markers (Fig. 1). We applied two different acceptance criteria to the data with different stringency regimes. One more naive one that accepted any two positives out of the twelve *PCR replicates* (from now on referred to as lax), and one stringent one that only accepted taxonomic assignments that were positively detected in both *extraction replicates* (*A & B*, Fig. 3). Our lax approach refers to one of the approaches of Ficetola et al. [19] where they evaluated different statistical approaches developed to estimate occupancy in the presence of observational errors and has been applied in other studies (e.g. [13]). The reason for conservatively omitting assignments that appeared in only one *extraction replicate* was to rule out sample cross-contamination during DNA extraction. In addition, we only accepted assignments with ten or more reads per marker, if only one marker was sequenced. If a species was assigned in more than one marker (e.g. *12S* and *16S*), we accepted the assignment even if in one sequencing run the number of reads was below ten.

**Figure 3:**
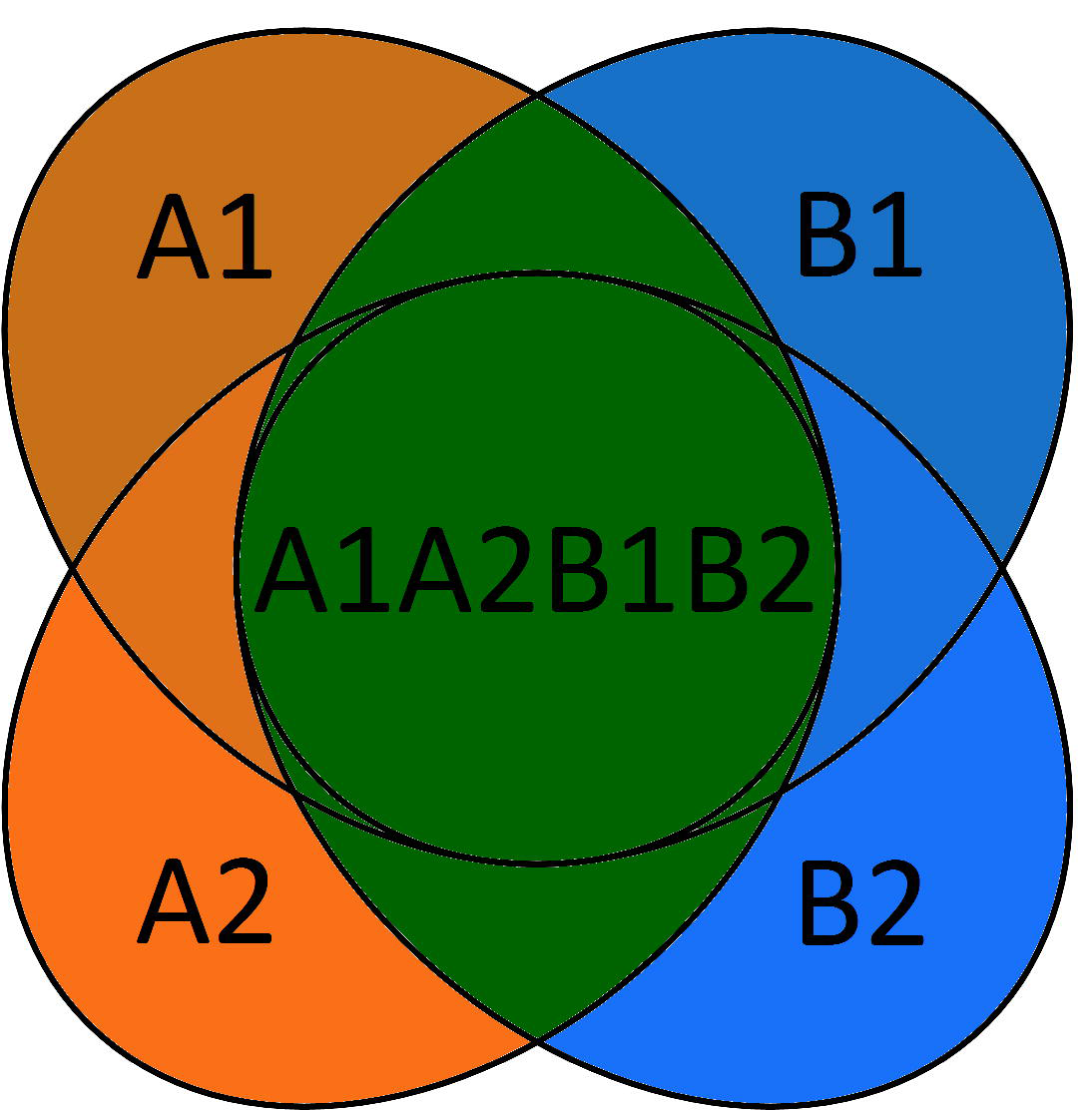
For the stringent acceptance criterion we only accepted taxonomic assignments that were positively detected in both *extraction replicates A* and *B* (green colour). The numbers (1 & 2) refer to the two *PCR replicates* for each extraction replicate.

Due to the imperfect PCR amplification of markers (the small *16S* fragment amplified better than the longer *CytB* fragment) and missing reference sequences in the database or shared sequence motifs between species, reads sometimes were assigned to species level for one marker but only to genus level for another marker. Thus, the final identification of species could not be automated, and manual inspection and curation was needed. For each assignment, three parameters were taken into consideration: number of sequencing reads, the mean probability estimate derived from *PROTAX*, and the mean sequence similarity to the reference sequences based on LAST.

#### Shot-gun sequencing to quantify mammalian DNA content

As the success of the metabarcoding largely depends on the mammal DNA quantity in our leech bulk samples we quantified the mammalian DNA content in a subset of 58 of our leech samples using shotgun sequencing. Extracted DNA was sheared with a Covaris M220 focused-ultra-sonicator to a peak target size of 100-200 bp, and re-checked for size distribution. Double-stranded Illumina sequencing libraries were prepared according to a ligation protocol designed by Fortes and Paijmans [51] with single 8 nt indices. All libraries were pooled equimolarly and sequenced on the MiSeq using the v3 150-cycle kit. We demultiplexed reads using bcl2fastq and cutadapt for trimming the adapters. We used BLAST search to identify reads and applied Metagenome Analyzer MEGAN [30] to explore the taxonomic content of the data based on the NCBI taxonomy. Finally we used KRONA [52] for visualisation of the results.

## Findings & Discussion

### Database curation

The Midori UNIQUE database (December 2015 version) contains 1,019,391 sequences across the four mitochondrial loci of interest (*12S*: 66,937; *16S*: 146,164; *CytB*: 223,247; COI: 583,043), covering all Metazoa. Of these, 258,225 (25.3%) derive from the four tetrapod classes (Amphibia: 55,254; Aves: 51,096; Mammalia: 101,106; Reptilia: 50,769). The distribution of these sequences between classes and loci, and the losses at each curation step are shown in Figure 4. In three of the four classes, there is a clear bias towards *CytB* sequences, with over 50% of sequences derived from this locus. In both Aves and Mammalia, the *16S* and *12S* loci are severely underrepresented at less than 10% each, while for Reptilia, *COI* is the least sequenced locus in the database.

**Figure 4:**
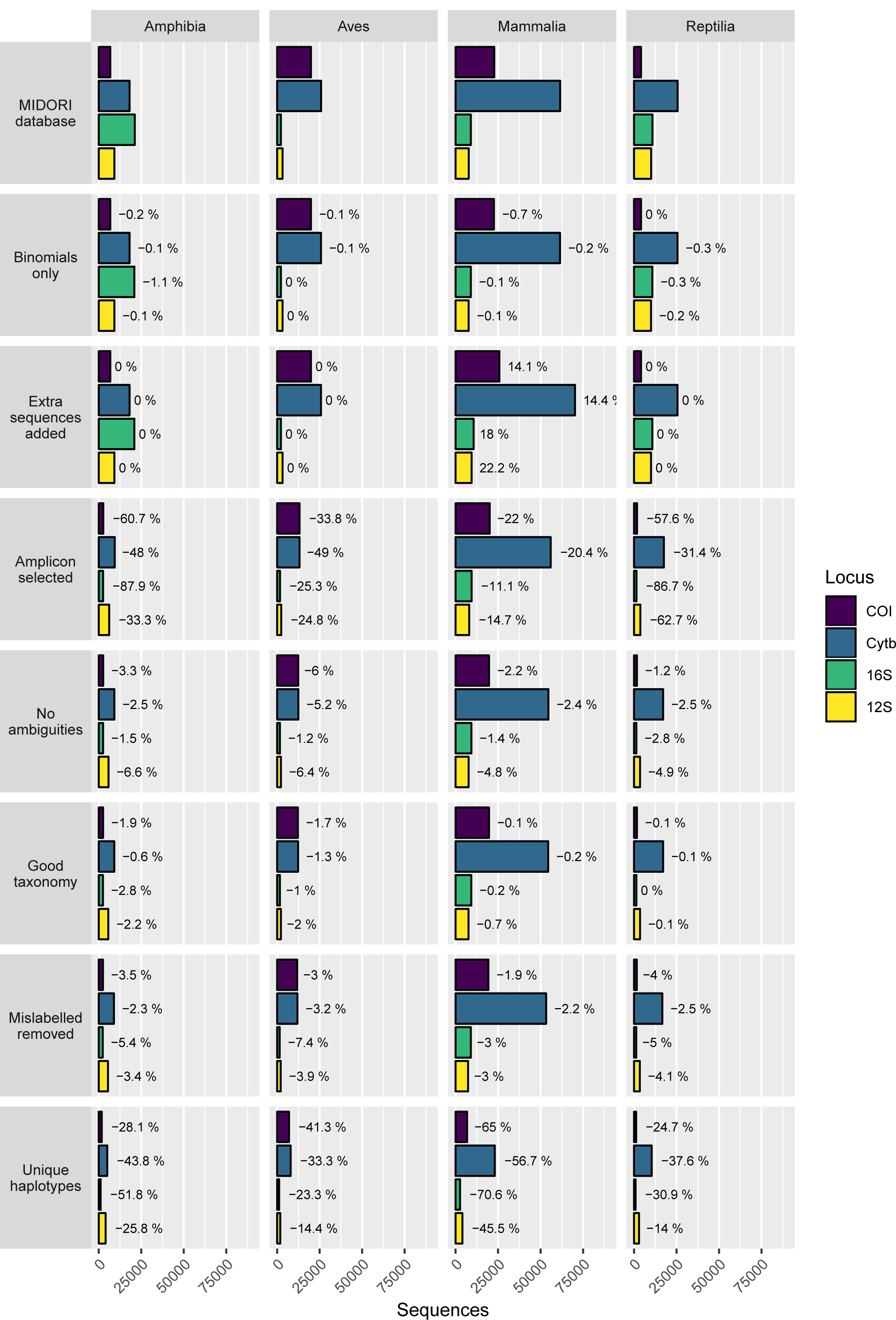
Data availability and percentage loss at each major step in the database curation procedure for each target amplicon and class of Tetrapoda. The number of sequences decreases between steps except “Extra sequences added” where additional target sequences are included for Mammalia and there is no change for the other three classes.

The numbers of sequences and rates of loss due to our curation steps varied among taxonomic classes and the four loci, although losses were observed between steps in almost all instances. The most significant losses followed amplicon trimming and removal of non-unique sequences. Amplicon trimming led to especially high losses in Amphibia and *16S*, indicating that data published on GenBank for this class and marker do not generally overlap with our amplicons. Meanwhile, the high level of redundancy in public databases was highlighted by the significant reduction in the number of sequences during the final step of removing redundant sequences – in all cases over 10% of sequences was discarded, with some losses exceeding 50% (Mammalia: *COI*, *CytB*, *16S*; Amphibia: *16S*).

Data loss due to apparent mislabelling ranged between 1.9% and 7.4% and was thus generally higher than similar estimates for curated microbial databases [29]. SATIVA flags potential mislabels and suggests an alternative label supported by the phylogenetic placement of the sequences, allowing the user to make an appropriate decision on a case by case basis. The pipeline pauses after this step to allow such manual inspection to take place. However, for the current database, the number of sequences flagged was large (4378 in total), and the required taxonomic expertise was lacking, so all flagged sequences from non-target species were discarded to be conservative. The majority of mislabels were identified at species level (3053), but there were also significant numbers at genus (788), family (364) and order (102) level. Two to three sequences from Bornean mammal species were unflagged in each amplicon to retain the sequences in the database. This was important as in each case these were the only reference sequences available for the species. Additionally, *Muntiacus vaginalis* sequences that were automatically synonymised to *M. muntjak* based on the available information in the Catalogue of Life were revised back to their original identifications to reflect current taxonomic knowledge.

### Database composition

The final database was skewed even more strongly towards *CytB* than was the raw database. It was the most abundant locus for each class and represented over 60% of sequences for both Mammalia and Reptilia. In all classes, *16S* made up less than 10% of the final database, with Reptilia COI also at less than 10%.

Figure 5 shows that most species represented in the curated database for any locus have just one unique haplotype against which HTS reads can be compared; only a few species have many haplotypes. The prevalence of species with 20 or more haplotypes is particularly notable in *CytB* where the four classes have between 25 (Aves) and 265 (Mammalia) species in this category. The coloured circles in Figure 5 also show that the species of the taxonomy are incompletely represented across all loci, and that coverage varies significantly between taxonomic groups. In spite of global initiatives to generate *COI* sequences [53], this marker does not offer the best species-level coverage in any class and is a poor choice for Amphibia and Reptilia (<15% of species included). Even the best performing marker, *CytB*, is not a universally appropriate choice, as Amphibia is better covered by *12S*. These differences in underlying database composition will impact the likelihood of obtaining accurate taxonomic assignment for any one species from any single marker. Further barcoding campaigns are clearly needed to fill gaps in the reference databases for all markers and all classes to increase the power of future e/iDNA studies. As the costs of HTS decrease, we expect that such gap-filling will increasingly shift towards sequencing of whole mitochondrial genomes of specimen obtained from museum collections, trapping campaigns etc. [34], reducing the effect of marker choice on detection likelihood. In the meantime, however, the total number of species covered by the database can be increased by combining multiple loci (here, up to four) and thus the impacts of database gaps on correctly detecting species can be minimized ([54]; Fig. 6).

**Figure 5:**
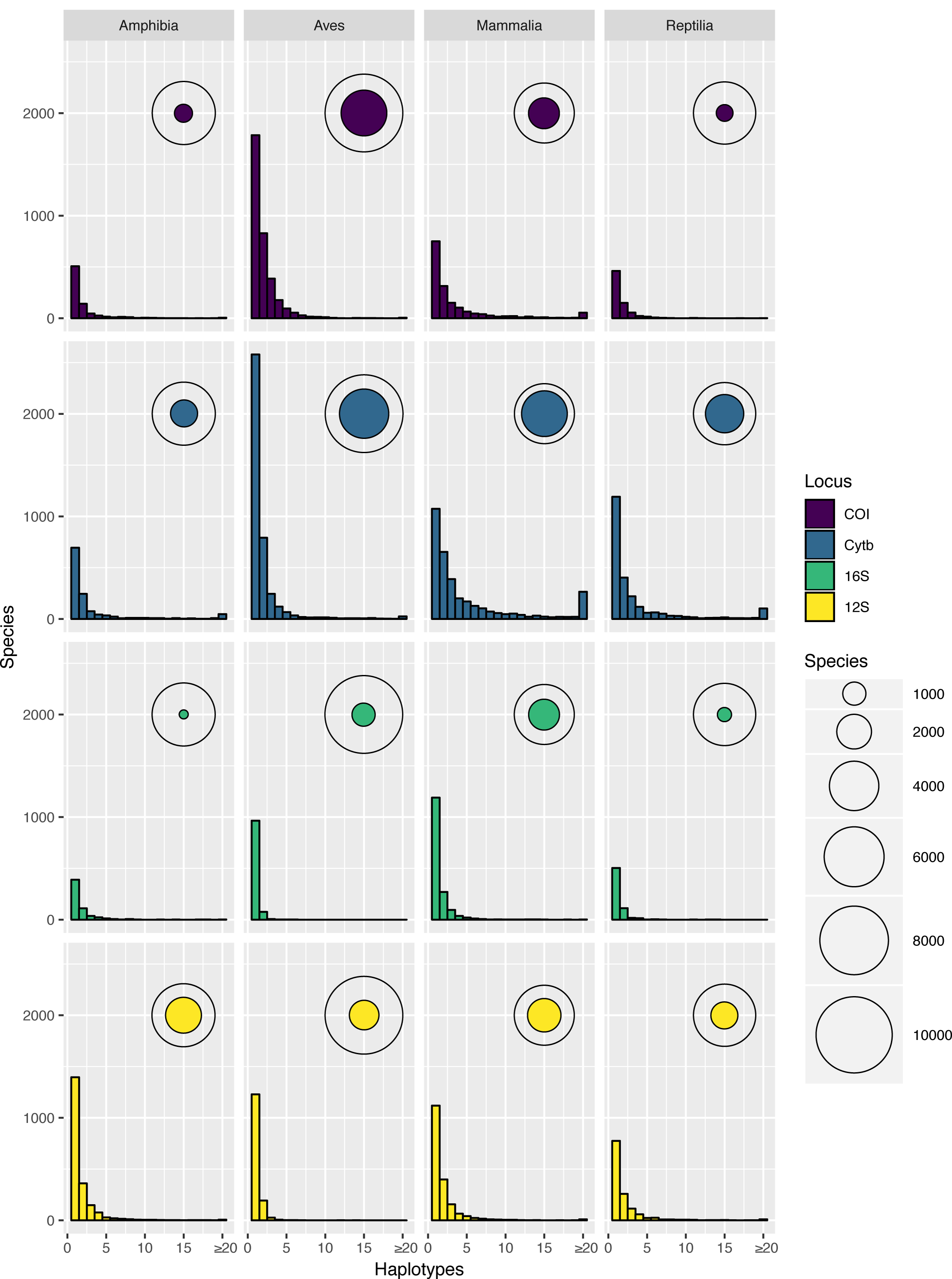
Haplotype number by species (frequency distribution) and the total number of species with at least one haplotype, shown relative to the total number of species in the taxonomy for that category (bubbles), shown for each marker and class of Tetrapoda. The proportion of species covered by the database varies between categories but in all cases a majority of recovered species are represented by a single unique haplotype.

**Figure 6:**
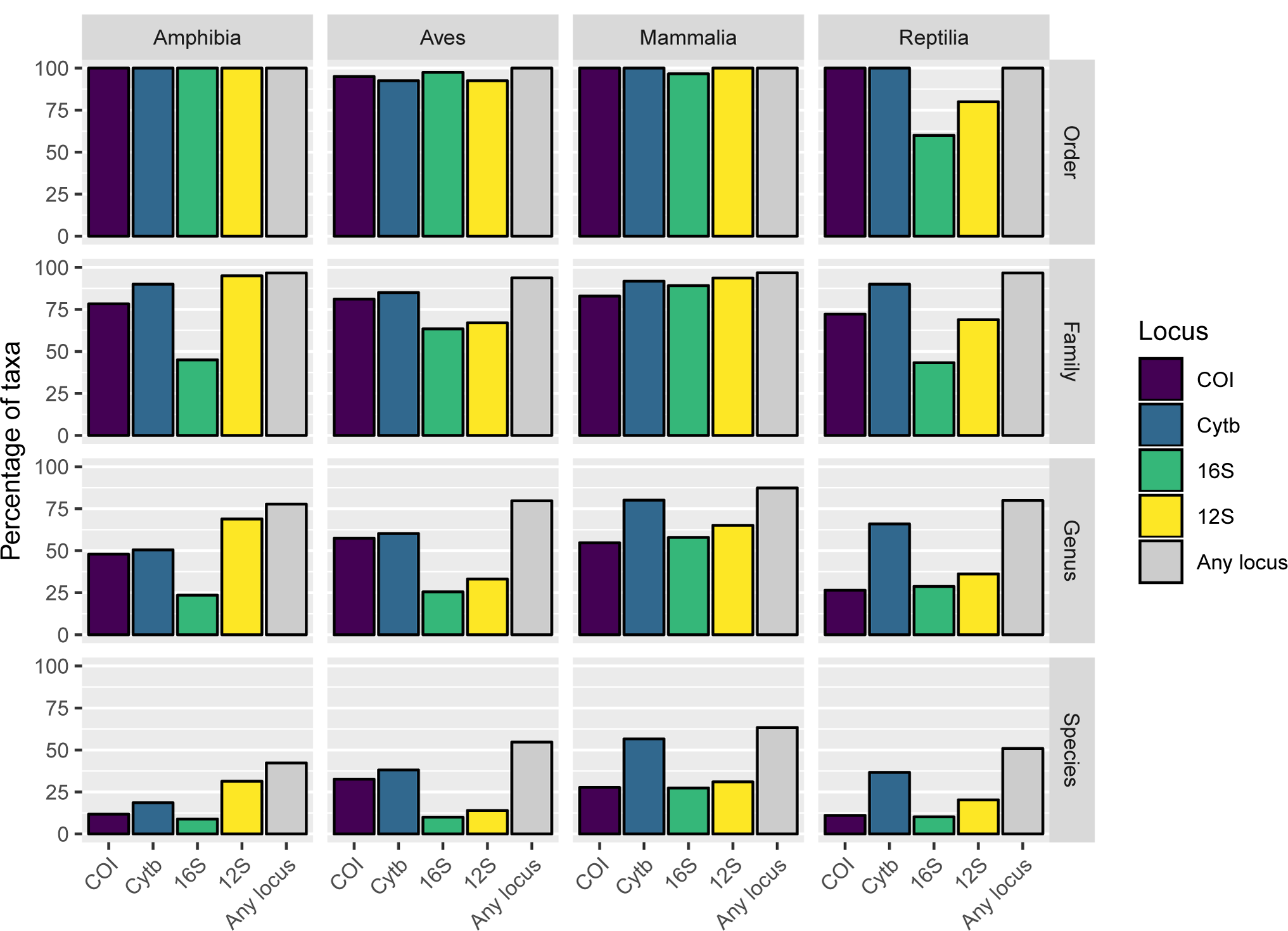
The percentage of the full taxonomy covered by the final database at each taxonomic level for each class of Tetrapoda. Includes the percentage of taxa represented by each marker and all markers combined. In all cases taking all four markers together increases the proportion of species, genera and families covered by the database, but it remains incomplete when compared with the full taxonomy.

In the present study, the primary target for iDNA sampling was the mammal fauna of Malaysian Borneo, and the 103 species expected in the sampling area represent an informative case study highlighting the deficiencies in existing databases (Fig. 7). Nine species are completely unrepresented while only slightly over half (55 species) have at least one sequence for all of the loci. Individually, each marker covers over half of the target species, but none achieves more than 85% coverage (*12S*: 75 species; *16S*: 68; *CytB*: 88; COI: 66). Equally striking is the lack of within-species diversity, as most of the incorporated species are represented by only a single haplotype per locus. Some of the species have large distribution ranges, so it is likely that in some cases the populations on Borneo differ genetically from the available reference sequences, possibly limiting assignment success. Only a few expected species have been sequenced extensively, and most are of economic importance to humans (e.g. *Bos taurus, Bubalus bubalis, Macaca spp, Paradoxurus hermaphroditus, Rattus spp., Sus scrofa*), with as many as 100 haplotypes available (*Canis lupus*). Other well-represented species (≥20 haplotypes) present in the sampling area include several Muridae (*Chiropodomys gliroides, Leopoldamys sabanus, Maxomys surifer, Maxomys whiteheadi*) and the leopard cat (*Prionailurus bengalensis*).

**Figure 7:**
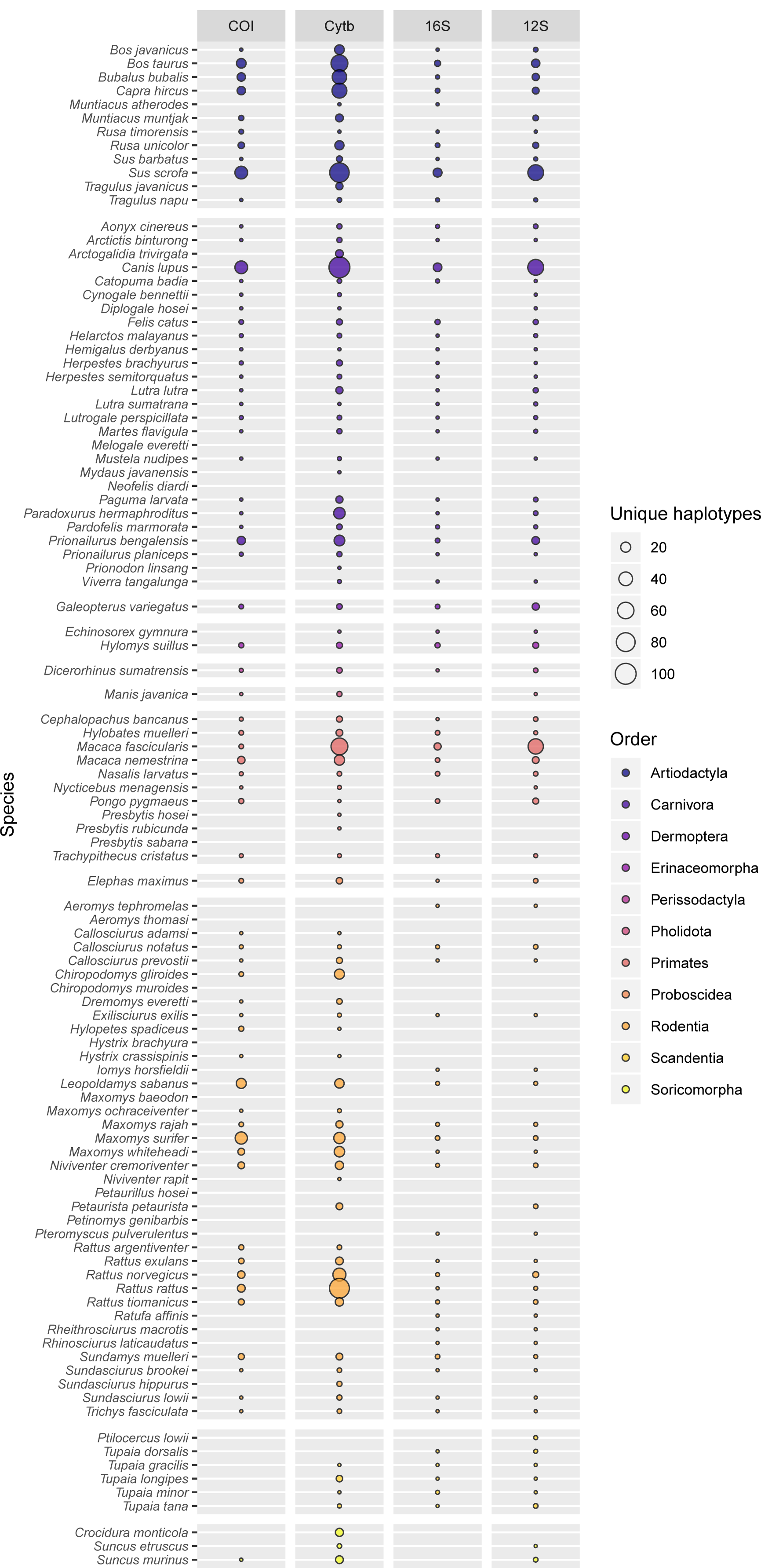
The number of unique haplotypes per marker for each of the 103 mammal species expected in the study area. Bubble size is proportional to the number of haplotypes and varies between 0 and 100. Only 55 species have at least one sequence per marker and nine species are completely unrepresented in the current database.

### Laboratory workflow

Shotgun sequencing of a subset of our samples revealed that the median mammalian DNA content was only 0.9%, ranging from 0% to 98%. These estimates are approximate, but with more than 75% of the samples being below 5%, this shows clearly the scarcity of target DNA in bulk iDNA samples. The generally low DNA content and the fact that the target DNA is often degraded make enrichment of the target barcoding loci necessary. We used PCR with high cycle numbers to obtain enough DNA for sequencing. However, this second step increases the risk of PCR error: artificial sequence variation, non-target amplification, and/or raising contaminations up to a detectable level.

We addressed these problems by running two *extraction replicates*, two *PCR replicates*, and a multi-marker approach. The need for *PCR replicates* has been acknowledged and addressed extensively in ancient DNA studies [16] and has also been highlighted for metabarcoding studies [19; 20; 55; 56]. Despite this, many e/iDNA studies do not carry out multiple *PCR replicates* to detect and omit potential false sequences. In addition, *extraction replicates* are seldom applied, despite the evidence that cross-sample DNA contamination can occur during DNA extraction [57; 58; 59]. We only accepted sequences that appeared in a minimum of two independent PCRs for the lax and for the stringent criterion, where it has to occur in each *extraction replicate A* and *B* (Fig. 1). The latter acceptance criterion is quite conservative and produces higher false negative rates than e.g. accepting occurrence of at least two positives. However, it also reduces the risk of accepting a false positives compared to it (see Supplemental Fig. 1. for a simulation of false positive and false negatives rates within a PCR), especially with increasing risk of false positive occurrence in a PCR for e.g. example due to higher risk of contamination etc.. Metabarcoding studies are very prone to false negatives, and downstream analyses like occupancy models for species distributions can account for imperfect detection and false negatives. However, methods for discounting false positive detections are not well developed [60]. Thus we think it is more important to avoid false positives, especially if the results will be used to make management decisions regarding rare or endangered species. In contrast, it might be acceptable to use a relaxed acceptance criterion for more common species, as long as the rate false-positives/true-positives is small and does not affect species distribution estimates. Employing both of our tested criteria researchers could flag unreliable assignments and management decisions can still use this information, but now in a forewarned way. An alternative to our acceptance criteria could be use the PCR replicates itself to model the detection probability within a sample using an occupancy framework [20; 60; 61; 62].

We used three different loci to correct for potential PCR-amplification biases. We were, however, unable to quantify this bias in this study due to the high degradation of the target mammalian DNA, which resulted in much higher overall amplification rates for *16S*, the shortest of our PCR amplicons. For *16S*, 85% of the samples amplified, whereas for *CytB* and *12S*, only 57% and 44% amplified, respectively. Also the read losses due to trimming and quality filtering were significantly lower for the *16S* sequencing runs (1.3% and 5.3% in average, Supplemental Table 3) compared to the sequencing runs for the longer fragments of *12S* and *CytB* (65.3% and 44.3% in average, Supplemental Table 3). Despite the greater taxonomic resolution of the longer *12S* and *CytB* fragments, our poorer amplification and sequencing results for these longer fragments emphasize that e/iDNA studies should generally focus on short PCR fragments to increase the likelihood of positive amplifications of the degraded target DNA. In the case of mammal-focussed e/iDNA studies, developing a shorter (100 bp) *CytB* fragment would likely be very useful.

Another major precaution was the use of twin-tagging for both PCRs (Fig. 2). This ensures that unlabelled PCR products are never produced and allows us to multiplex a large number of samples on a single run of Illumina MiSeq run. Just 24 sample *tags 1* and 20 plate *tags 2* allow the differentiation of up to 480 samples with matching tags on both ends. The same number of individual primers would have needed longer tags to maintain enough distance between them and would have resulted in an even longer adapter-tag overhang compared to primer length. This would have most likely resulted in lower binding efficiencies due to steric hindrances of the primers. Furthermore, this would have resulted in increased primer costs. Thus our approach reduced sequencing and primer purchase costs while at the same time largely eliminating sample mis-assignment via tag jumping, because tag-jump sequences have non-matching forward and reverse *tag 1* sequences [43]. We estimated the rate of tag jumps producing non-matching *tag 1* sequences to be 1 to 5%, and these were removed from the dataset (Table 4). For our sequenced PCR plates, the rate of non-matching *tag 2* tags was 2%. These numbers are smaller than data from Zepeda-Mendoza et al. [56] who reported on sequence losses of 19% to 23% due to unused tag combinations when they tested their DAMe pipeline to different datasets built using standard blunt-end ligation technique. Although their numbers might not be one-to-one comparable to our results as they counted unique sequences, and we report on read numbers, our PCR libraries with matching barcodes seem reduce the risk of tag jumping compared to blunt-end ligation techniques. For the second PCR round, we used the same tag pair *tag 2* for all 24 samples of a PCR plate. In order to reduce cost we tested pooling these 24 samples prior to the second PCR round, but we detected a very high tag jumping rate of over 40% (Table 4), which ultimately would increase cost through reduced sequencing efficiency. Twin-tagging increases costs because of the need to purchase a larger number of primer pairs but at the same time it increases confidence in the results.

**Table 4:** Number of reads per sequencing run and the numbers of reads with matching, non-matching or unidentifiable tags for seven of the eight sequencing runs^*^.

|  | total<br>reads | matching<br>tag 2<br>reads | non-matching<br>tag 2<br>reads | %<sup>1</sup> | matching<br>tag 1<br>reads | non-matching<br>tag 1<br>reads | %<sup>2</sup> | erroneous<br>tag 1<br>reads | %<sup>2</sup> |
| --- | --- | --- | --- | --- | --- | --- | --- | --- | --- |
| **SeqRun01** | 18,438,517 | 18,102,702 | 282,419 | 1.5 | 17,514,515 | 451,028 | 2.5 | 137,159 | 0.8 |
| **SeqRun02** | 25,385,558 | 24,596,380 | 626,245 | 2.5 | 23,426,084 | 612,045 | 2.5 | 558,251 | 2.3 |
| **SeqRun03** | 14,875,796 | 14,393,884 | 343,528 | 2.3 | 13,766,187 | 426,181 | 3.0 | 201,516 | 1.4 |
| **SeqRun04** | 2,027,794 | 1,935,149 | 56,077 | 2.8 | 1,806,655 | 88,307 | 4.6 | 40,187 | 2.1 |
| **SeqRun05** | 18,221,504 | 17,500,366 | 421,588 | 2.3 | 16,793,851 | 482,365 | 2.8 | 161,458 | 0.9 |
| **SeqRun06** | 20,718,202 | 19,874,913 | 429,048 | 2.1 | 19,317,305 | 371,048 | 1.9 | 81,422 | 0.4 |
| **SeqRun07** | 24,604,610 | 23,746,938 | 663,730 | 2.7 | 22,446,187 | 497,366 | 2.1 | 803,385 | 3.4 |
| **Total** | 124,271,981 | 120,150,332 | 2,822,635 | 2.3 | 115,070,784 | 2,928,340 | 2.5 | 1,983,378 | 1.7 |
| **IndexRun** | 10,276,093 | 10,116,808 | NA | NA | 5,841,190 | 4,186,688 | 41.4 | 88,930 | 0.9 |

<sup>1</sup> refers to total reads
<sup>2</sup> refers to matching tag 2
\*Sequencing run SeqRun08 run contained libraries of another project, thus we were unable to provide a number of raw reads.

Tagging primers in the first PCR reduces the risk of cross-contamination via aerosolised PCR products. However, we would not be able to detect a contamination prior the second PCR from one plate to another, as we used the same 24 tags (*tag 1*) for all plates. Nevertheless such a contamination is very unlikely to result in any accepted false positive as it would be improbable to contaminate both the A and B replicates, given the exchange of all reagents and the time gap between the PCRs. Previous studies have shown that unlabelled volatile PCR products pose a great risk of false detections [63], a risk that is greatly increased if a high number of samples are analysed in the laboratories [13]. Also, in laboratories where other research projects are conducted, this approach allows the detection of cross-experiment contamination. Therefore, we see a clear advantage of our approach over ligation techniques when it comes to producing sequencing libraries, as the Illumina tags are only added after the first PCR, and thus the risk of cross contamination with unlabelled PCR amplicons is very low.

### Assignment results

A robust assignment of species is an important factor in metabarcoding as an incorrect identification might result incorrect management interventions. The reliability of taxonomic assignments is expected to vary with respect to both marker information content and database completeness, and this is reflected in the probability estimates provided by *PROTAX*. In a recent study, less than 10% of the mammal assignments made at species level against a worldwide reference database were considered reliable with the short *16S* amplicon, but this increased to 46% with full-length *16S* sequences [31]. In contrast, in the same study over 80% of insect assignments at species level were considered reliable with a more complete, geographically restricted database of full-length COI barcodes. A similar pattern was observed in our data during manual curation of the assignment results – there was more ambiguity in the results for the short *16S* amplicon than for other markers. However, due to the limited amount of often degraded target DNA in e/iDNA samples, short amplicons amplify much better. In our case, this had the drawback that some species lacked any interspecific variation, and thus sequencing reads shared 99%-100% identity for several species. For example, our only *16S* reference of *Sus barbatus* was 100% identical to *S. scrofa*. But as latter species does not occur in the studied area we could assign all reads manually to *S. barbatus*. In several cases we were able to confirm *S. barbatus* by additional *CytB* results, highlighting the usefulness of multiple markers.

Another advantage of multiple markers is the opportunity to fill gaps in the reference database. For example, we lacked *16S* reference sequences for *Hystrix brachyura*, and reads were assigned by *PROTAX* only to the unknown species *Hystrix sp*.. In one sample, however, almost 5000 *CytB* reads could be confidently assigned to *Hystrix brachyura*, and thus we used the Hystrix sp. *16S* sequences in the same sample to build a consensus *16S* reference sequence for *Hystrix brachyura* for future analyses. In another example we had *CytB* reads assigned to *Mydaus javanicus*, the Sunda stink-badger in one sample but *12S* reads assigned to *Mydaus sp*. in another one. As we lacked a *12S Mydaus* reference and as there is only one *Mydaus* species on Borneo we could assume that this second sample is most likely also *Mydaus javanicus*.

We also inferred that PCR and sequencing errors resulted in reads being assigned to sister taxa. We observed that a high number of reads of a true sequence were assigned to a species and a lower number of noise sequences were assigned to a sister taxon. Such a pattern was observed for ungulates, especially deer that showed little variance in *16S*. It is hard to identify and control for such pattern automatically, and it highlights the importance of visual inspection of the results.

For the more lax criterion (two positive *PCR replicates*) we accepted 190 species assignments out of 109 leech samples. Under the stringent criterion (i.e. having positive detections in both *extraction replicates A* and *B*) we accepted about 14% assignments less; in total 162 vertebrate detections within 95 bulk samples (Table 5). For this genus we have two occurring species in the area. As the true occurrence of species within our leeches was unknown we cannot evaluate how many of the additional 27 detections in the lax criterion are false positives and how many might be false negatives for the stricter criterion. However, by accepting only positive *AB* assignment results, we increase the confidence of species detection, even if the total number of reads for that species was low. When it comes to rare or threated species this outweighs the risk of reporting false positives to our opinion. 48% of the assignments with the stringent criterion were present in all four *A1*, *A2*, *B1* and *B2*. 35% were present in at least three replicates (e.g. A1, A2, B1).

**Table 5:**
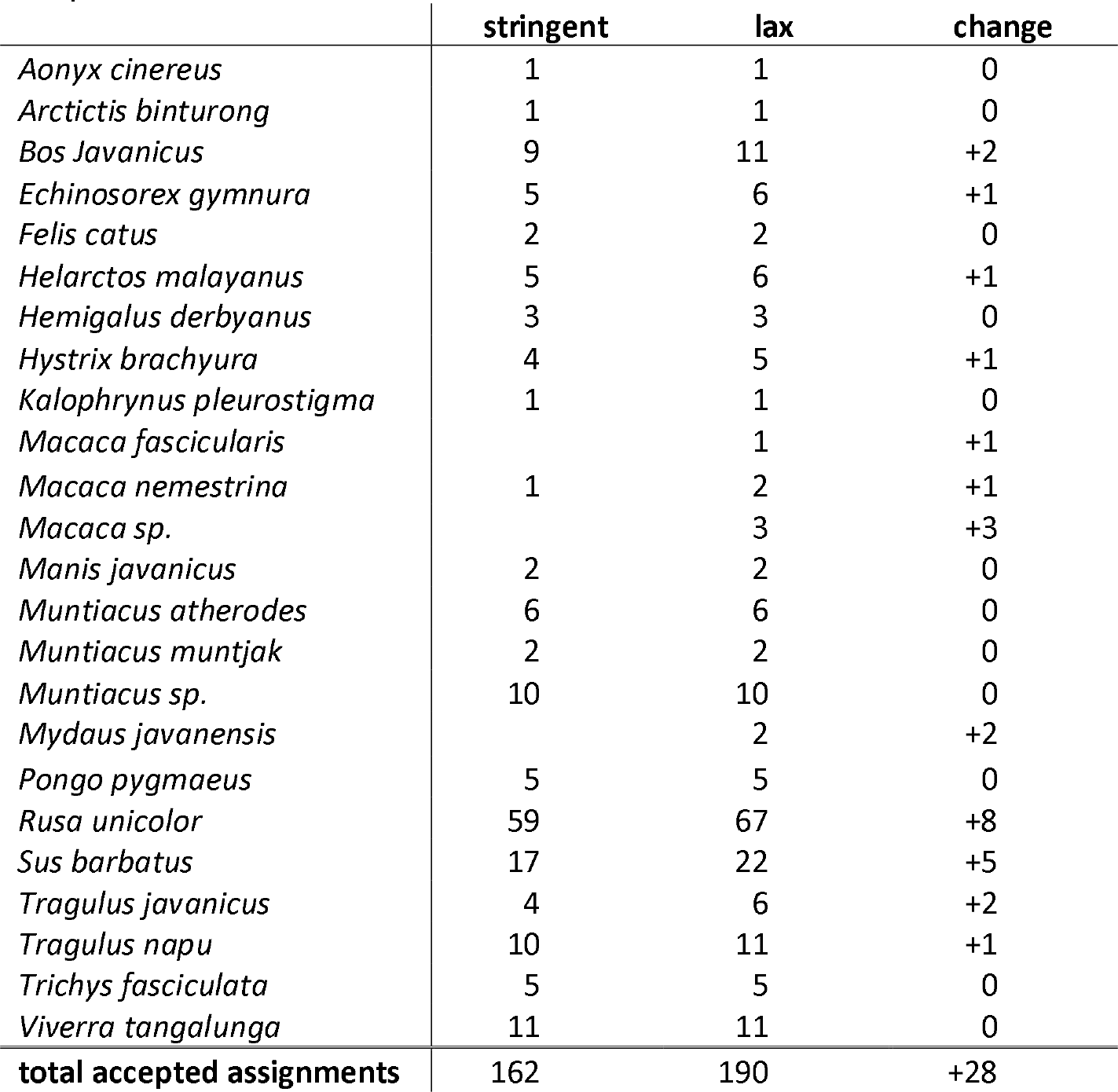
Number of accepted species assignments with two different acceptance criteria the more stringent criterion accepting only assignments occurring in both *extraction replicates* (A & B), and the more lax criterion accepting assignment two or more positives in any of the twelve *PCR replicates*.

The mean number of reads per sample used for the taxomomic assignment varied from 162,487 *16S* reads for SeqRun01 to 7,638 *CytB* reads for SeqRun05 (Supplemental Table 4). In almost all cases, however, the number of reads of an accepted assignment was high (median= 52,386; mean= 300,996; SD= 326,883). PCR stochasticity, primer biases, multiple species in individual samples, and pooling of samples exert too many uncertainties that could bias the sequencing results [64; 65]. Thus we do not believe that raw read numbers are the most reliable indicators of tetrapod DNA quantity in iDNA samples. Replication of detection is inherently more reliable. In contrast to our expectation that higher cycle number might be necessary to amplify even the lowest amounts of target DNA, our data do not support this hypothesis. Although we observed an increase in positive PCRs for A2/B2 (the 40-cycle *PCR replicates*), the total number of accepted assignments in *A1/B1* and *A2/B2* samples did not differ. This indicates first that high PCR cycle numbers mainly increased the risk of false positives and second that our multiple precautions successfully minimized the acceptance of false detections.

## Conclusion

Metabarcoding of e/iDNA samples will certainly become a very valuable tool in assessing biodiversity, as it allows to detect species non-invasively without the need to capture and handle the animals [66] and because sampling effort can often be greatly reduced. However, the technical and analytical challenges linked to sample types (low quantity and quality DNA) and poor reference databases have so far been insufficiently recognized. In contrast to ancient DNA studies where standardized laboratory procedures and specialized bioinformatics pipelines have been established and are followed in most cases, there is limited methodological consensus in e/iDNA studies, which reduces rigour. In this study, we present a robust metabarcoding workflow for e/iDNA studies. We hope that the provided scripts and protocols facilitate further technical and analytical developments. The use of e/iDNA metabarcoding to study the rarest and most endangered species such as the Saola is exciting, but geneticists bear the heavy responsibility of providing correct answers to conservationists.

## Acknowledgements

All authors thank the German Federal Ministry of Education and Research (BMBF FKZ: 01LN1301A) and the Leibniz-IZW for funding this study. We also thank the Sabah Forestry Department, especially Johnny Kissing, Peter Lagan and Datuk Sam Mannan for supporting the fieldwork and the Sabah Biodiversity Council for providing research, collection and export permits for this work. We are grateful to John Mathai, Seth Timothy Wong for conducting the field work and collecting the leeches. We are also grateful to Jörns Fickel, head of the Department Evolutionary Genetics of the Leibniz-IZW for continuous support and collaboration. Furthermore we would like to thank Sebastian Wieser for lab-support, Dorina Lenz and Anke Schmidt for their help and fruitful discussions. C.C.Y. Xu was also supported by the MEME Erasmus Mundus Programme in Evolutionary Biology, and by the Groningen University Fund and the Marco Polo Fund from the University of Groningen. D.W. Yu and C.C.Y. Xu were supported by the National Natural Science Foundation of China (41661144002, 31670536, 31400470, 31500305), the Key Research Program of Frontier Sciences, CAS (QYZDY-SSW-SMC024), the Bureau of International Cooperation project (GJHZ1754), the Strategic Priority Research Program of the Chinese Academy of Sciences (XDA20050202, XDB31000000), the Ministry of Science and Technology of China (2012FY110800), and the State Key Laboratory of Genetic Resources and Evolution at the Kunming Institute of Zoology.

---

**Supplemental Figure 1:**
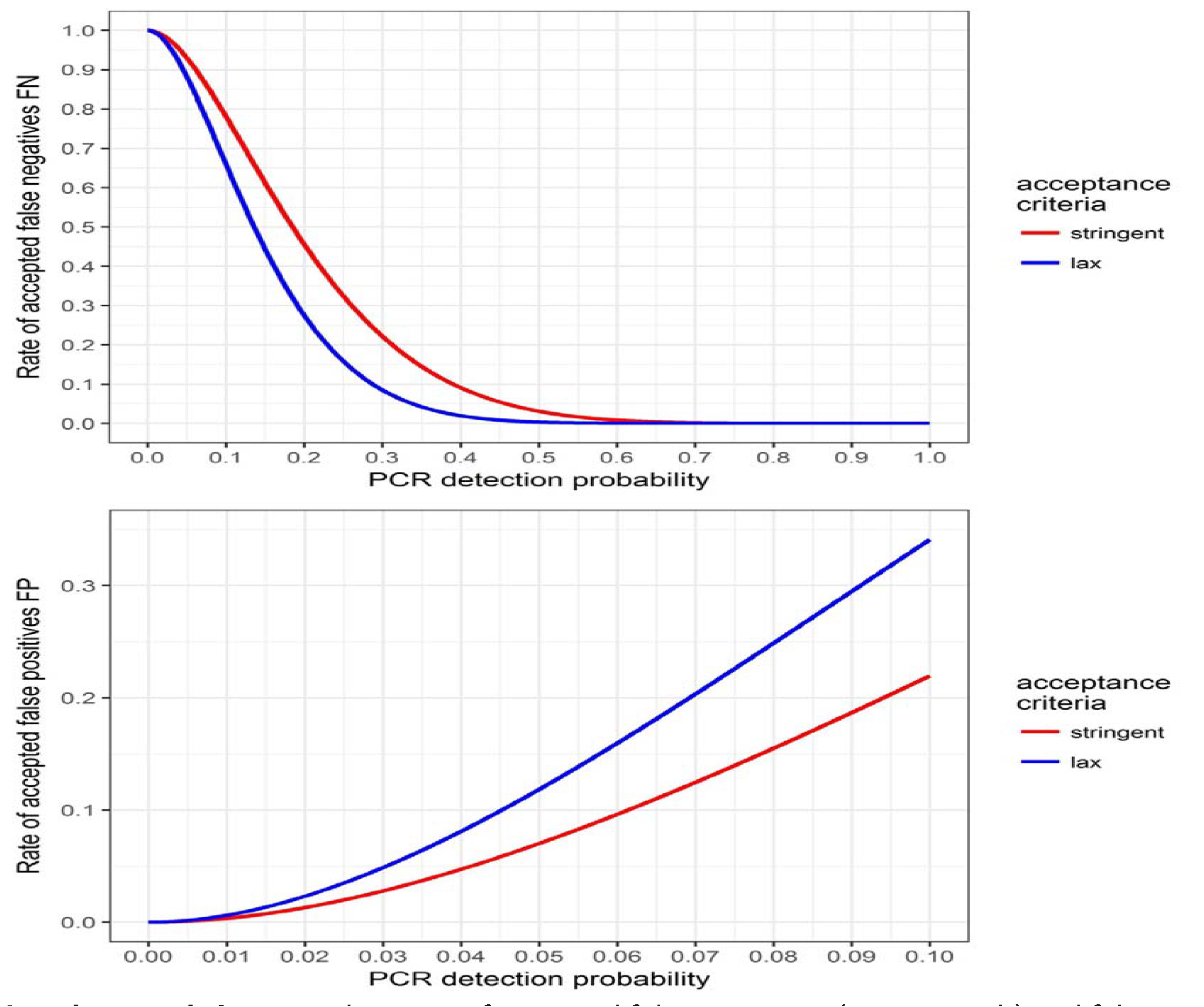
The rates of accepted false negatives (upper graph) and false positives (lower graph) for both our used acceptance criteria for varying PCR detection probabilities. The red line always denotes the stringent acceptance criterion that a positive is only accepted if it is present in at least one A and one B replicate. The lax criterion (blue) accepted at any two positives out of the twelve replicates. The stringent criterion poses a higher risk of accepting a false negative but it reduces clearly the risk of false positives, especially with increasing detection probability due to higher risk of contamination.

**Supplemental table 1:**
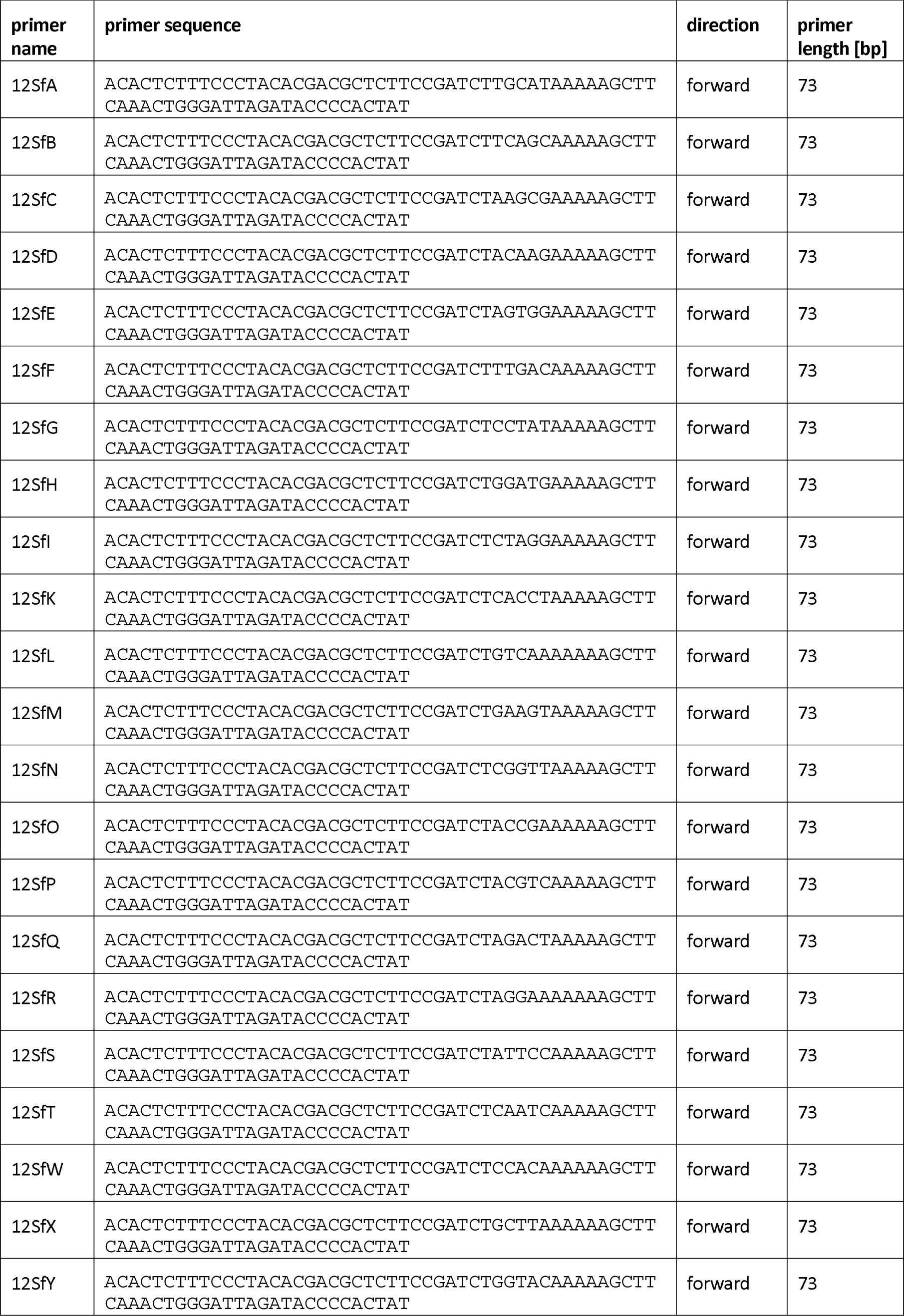
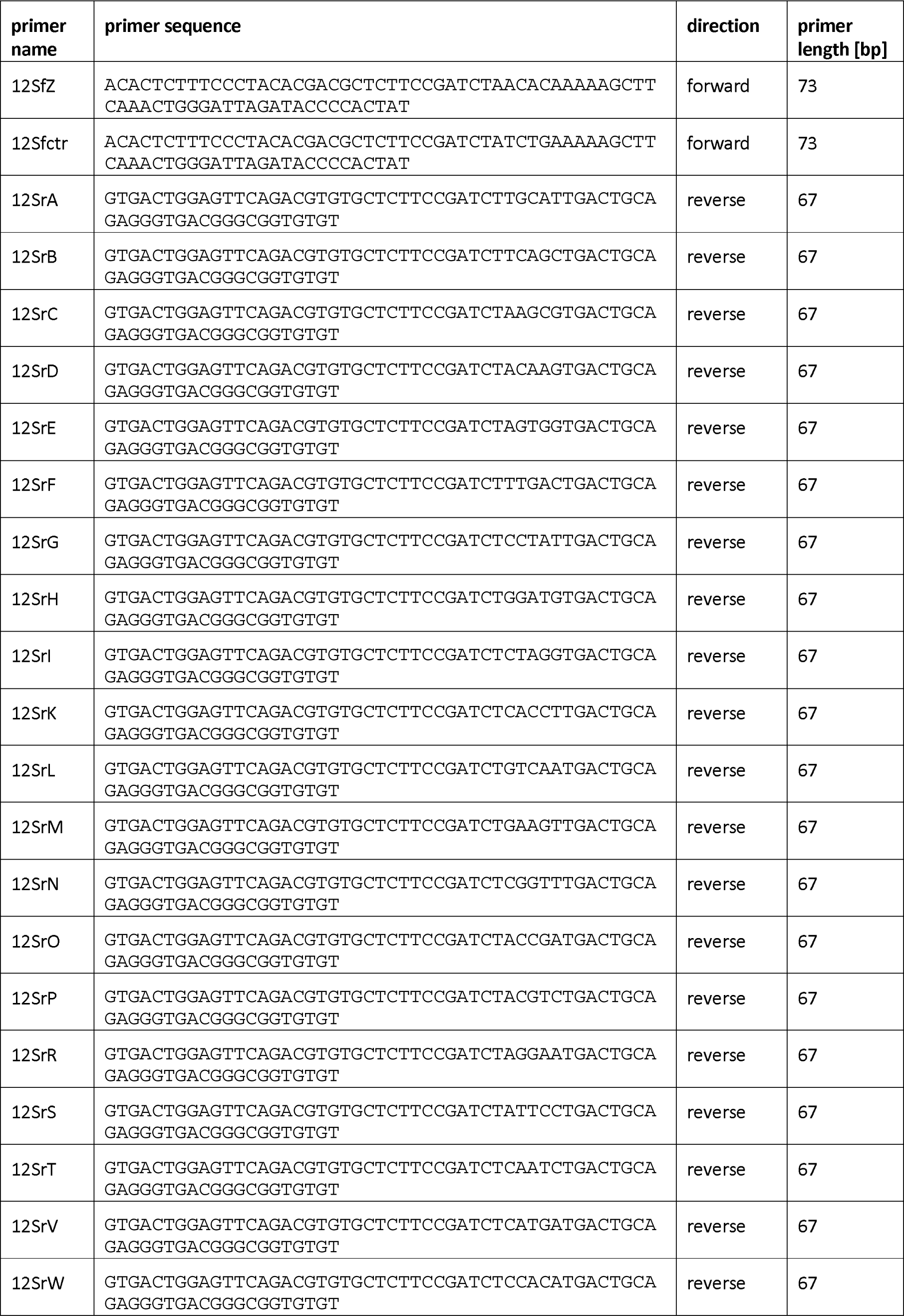
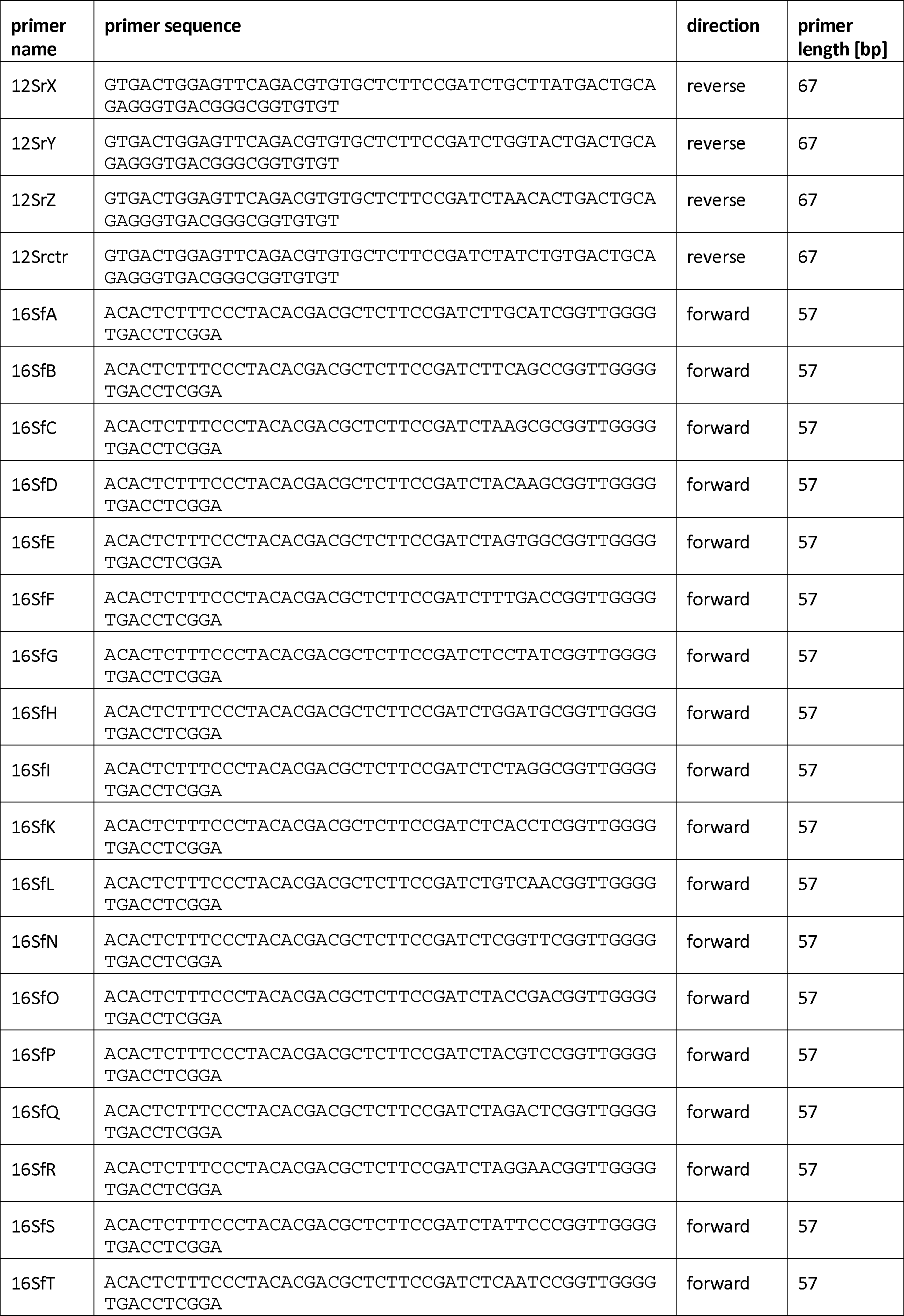
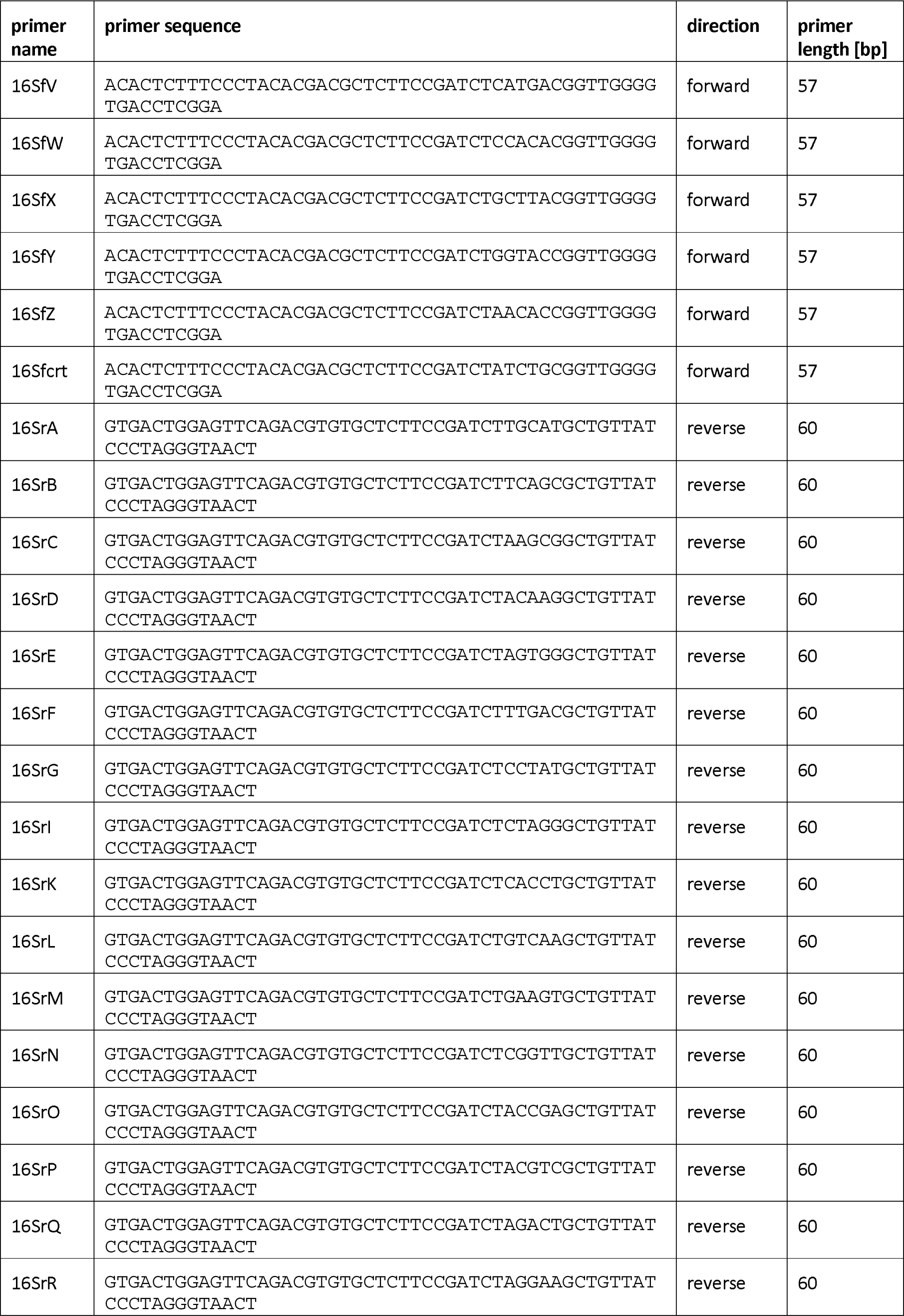
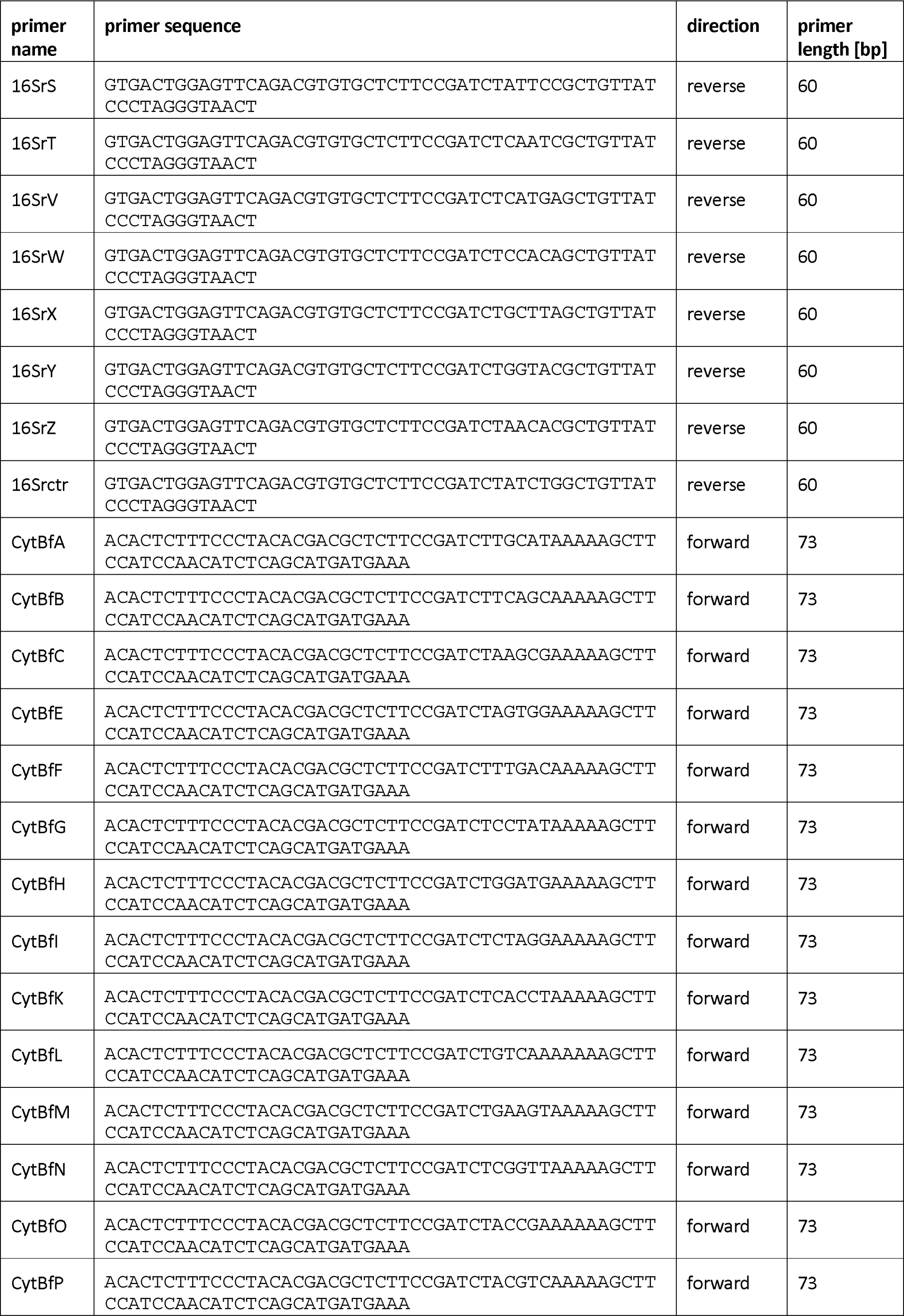
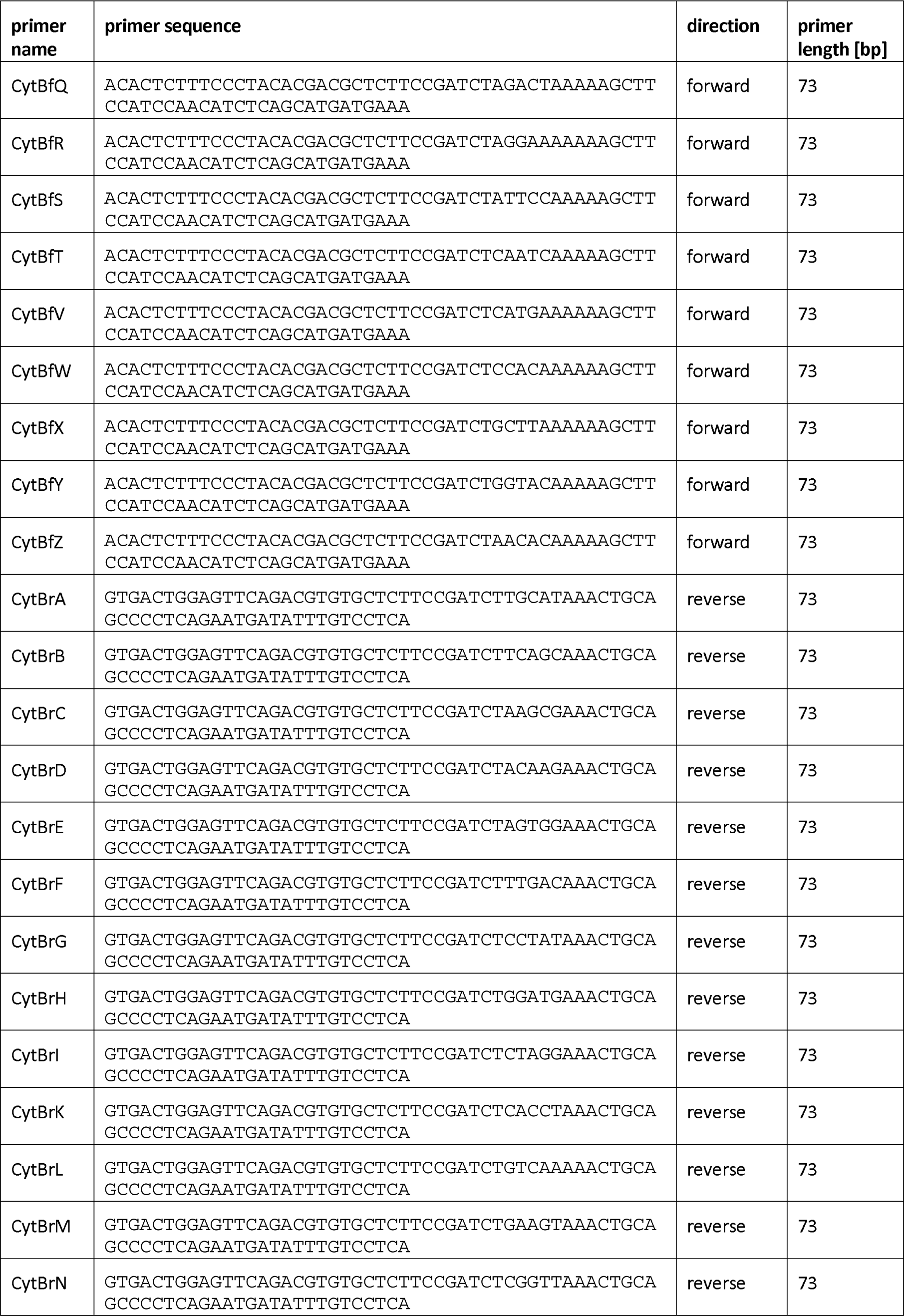
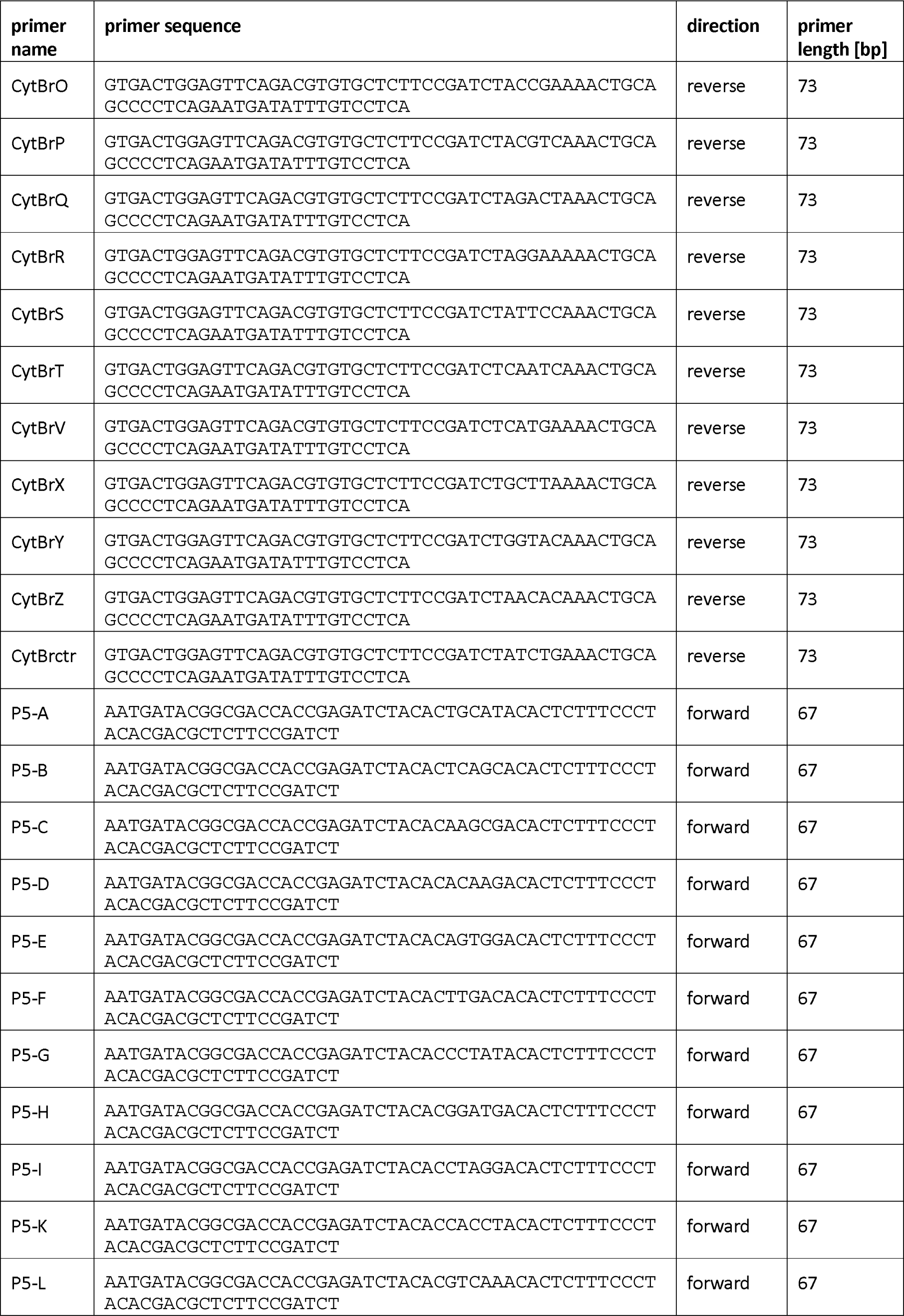
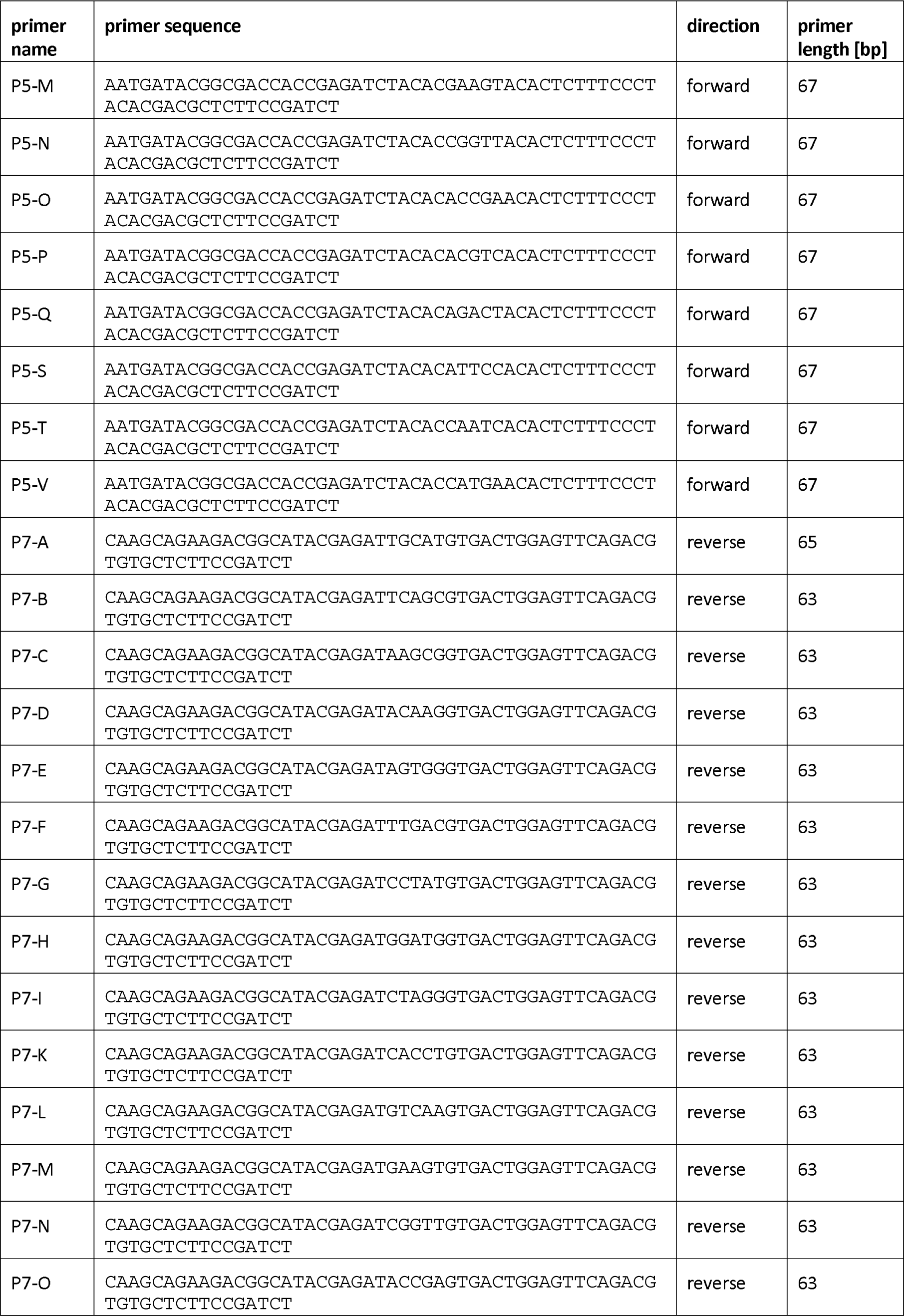
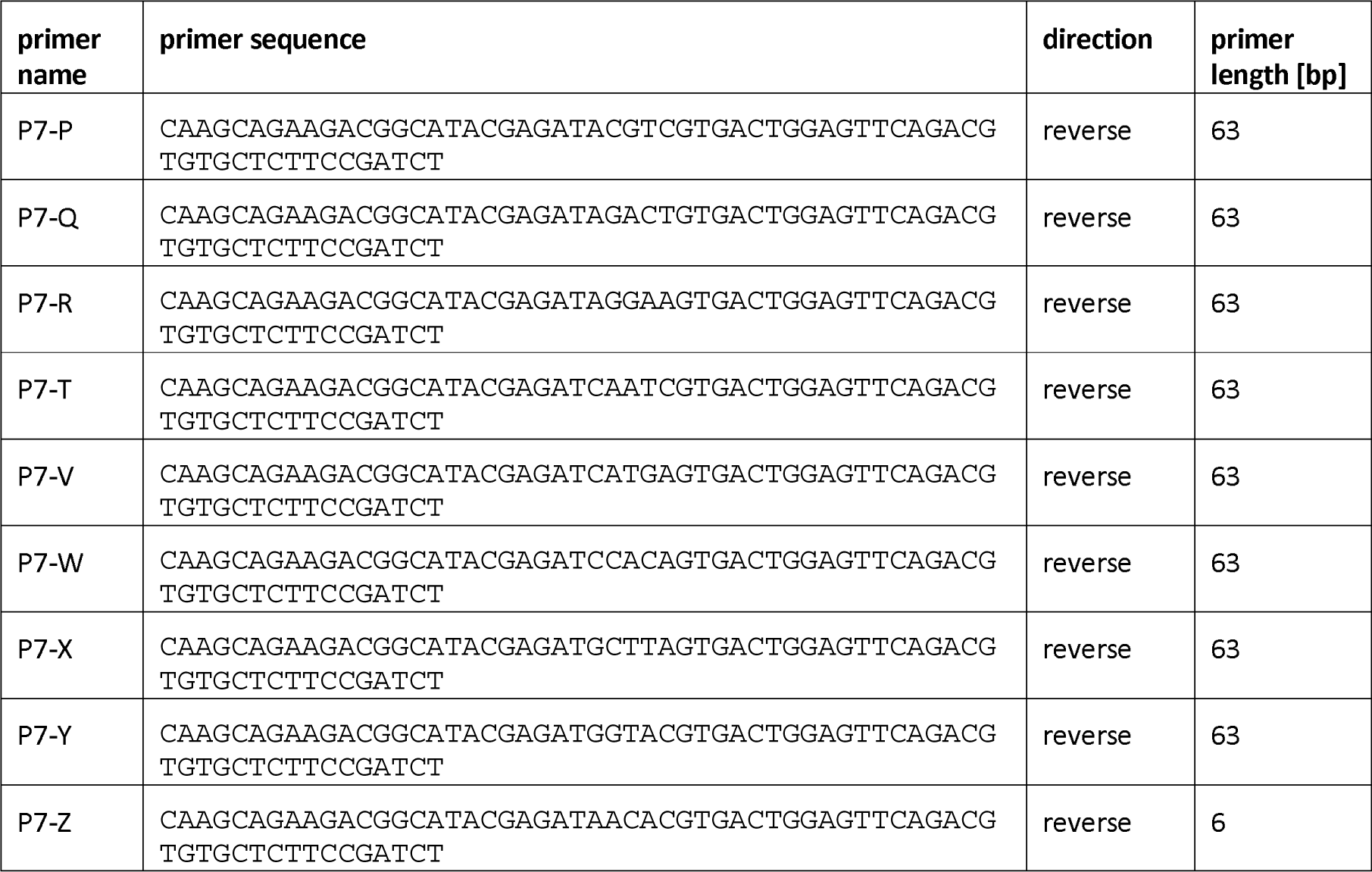
Complete list of all used primer sequences in 5’-3’ direction.

**Supplemental table 2:** List of Bornean species that were weighted in the *PROTAX* assignment.

| Species | Species | Species |
| --- | --- | --- |
| *Bos,javanicus* | *Arctictis,binturong* | *Chiropodomys,muroides* |
| *Bos,taurus* | *Arctogalidia,trivirgata* | *Leopoldamys,sabanus* |
| *Bubalus,bubalis* | *Cynogale,bennettii* | *Maxomys,baeodon* |
| *Capra,hircus* | *Diplogale,hosei* | *Maxomys,ochraceiventer* |
| *Muntiacus,atherodes* | *Hemigalus,derbyanus* | *Maxomys,rajah* |
| *Muntiacus,muntjak* | *Paguma,larvata* | *Maxomys,surifer* |
| *Rusa,timorensis* | *Paradoxurus,hermaphroditus* | *Maxomys,whiteheadi* |
| *Rusa,unicolor* | *Prionodon,linsang* | *Niviventer,cremoriventer* |
| *Sus,barbatus* | *Viverra,tangalunga* | *Niviventer,rapit* |
| *Sus,scrofa* | *Galeopterus,variegatus* | *Rattus,argentiventer* |
| *Tragulus,javanicus* | *Echinosorex,gymnura* | *Rattus,exulans* |
| *Tragulus,napu* | *Hylomys,suillus* | *Rattus,norvegicus* |
| *Canis,lupus* | *Dicerorhinus,sumatrensis* | *Rattus,rattus* |
| *Catopuma,badia* | *Manis,javanica* | *Rattus,tiomanicus* |
| *Felis,catus* | *Macaca,fascicularis* | *Sundamys,muelleri* |
| *Neofelis,diardi* | *Macaca,nemestrina* | *Aeromys,tephromelas* |
| *Pardofelis,marmorata* | *Nasalis,larvatus* | *Aeromys,thomasi* |
| *Prionailurus,bengalensis* | *Presbytis,hosei* | *Callosciurus,adamsi* |
| *Prionailurus,planiceps* | *Presbytis,rubicunda* | *Callosciurus,notatus* |
| *Herpestes,brachyurus* | *Presbytis,sabana* | *Callosciurus,prevostii* |
| *Herpestes,semitorquatus* | *Trachypithecus,cristatus* | *Dremomys,everetti* |
| *Mydaus,javanensis* | *Pongo,pygmaeus* | *Exilisciurus,exilis* |
| *Aonyx,cinereus* | *Hylobates,muelleri* | *Hylopetes,spadiceus* |
| *Lutra,lutra* | *Nycticebus,menagensis* | *Iomys,horsfieldii* |
| *Lutra,sumatrana* | *Cephalopachus,bancanus* | *Petaurillus,hosei* |
| *Lutrogale,perspicillata* | *Elephas,maximus* | *Petaurista,petaurista* |
| *Martes,flavigula* | *Hystrix,brachyura* | *Petinomys,genibarbis* |
| *Melogale,everetti* | *Hystrix,crassispinis* | *Pteromyscus,pulverulentus* |
| *Mustela,nudipes* | *Trichys,fasciculata* | *Ratufa,affinis* |
| *Helarctos,malayanus* | *Chiropodomys,gliroides* | *Rheithrosciurus,macrotis* |
| *Rhinosciurus,laticaudatus* | *Tupaia,dorsalis* | *Crocidura,monticola* |
| *Sundasciurus,brookei* | *Tupaia,gracilis* | *Suncus,etruscus* |
| *Sundasciurus,hippurus* | *Tupaia,longipes* | *Suncus,murinus* |
| *Sundasciurus,lowii* | *Tupaia,minor* |  |
| *Ptilocercus,lowii* | *Tupaia,tana* |  |

**Supplemental table 3:** Summary of the read losses of each sample during the read processing steps for each sequencing run seperately. The first line gives the raw read number per sample. The losses are given as percentage of each step; 1. merging of the R1/R2 reads of the Illumina sequencing done by *usearch* [43; 44], 2. clipping of primers and trimming of reads using *cutadapt* [45], 3. quality filtering and 4. dereplication, both using usearch.

|  | Step | Mean | SD | Median | Min | Max |
| --- | --- | --- | --- | --- | --- | --- |
| **SeqRun01** | raw | 72977 | 96466 | 74 | 1 | 422271 |
|  | merging | 7% | 11% | 2% | 1% | 50% |
|  | clipping & trimming | 2% | 14% | 0% | 0% | 100% |
|  | filtering | 4% | 11% | 2% | 1% | 100% |
| **SeqRun02** | raw | 97372 | 83870 | 117626 | 1 | 409999 |
|  | merging | 22% | 23% | 13% | 2% | 98% |
|  | clipping & trimming | 2% | 13% | 0% | 0% | 100% |
|  | filtering | 6% | 3% | 6% | 5% | 43% |
| **SeqRun03** | raw | 57359 | 123971 | 48 | 1 | 1105978 |
|  | merging | 5% | 3% | 5% | 1% | 11% |
|  | clipping & trimming | 43% | 40% | 28% | 0% | 100% |
|  | filtering | 37% | 20% | 29% | 24% | 100% |
| **SeqRun04** | raw | 8629 | 10184 | 2075 | 1 | 37592 |
|  | merging | 8% | 2% | 8% | 6% | 14% |
|  | clipping & trimming | 79% | 34% | 100% | 0% | 100% |
|  | filtering | 38% | 18% | 34% | 0% | 92% |
| **SeqRun05** | raw | 77936 | 193818 | 36 | 1 | 1081947 |
|  | merging | 34% | 17% | 36% | 4% | 89% |
|  | clipping & trimming | 50% | 41% | 59% | 0% | 100% |
|  | filtering | 53% | 19% | 51% | 0% | 100% |
| **SeqRun06** | raw | 80816 | 80656 | 87013 | 1 | 407872 |
|  | merging | 10% | 15% | 3% | 1% | 69% |
|  | clipping & trimming | 0% | 0% | 0% | 0% | 1% |
|  | filtering | 5% | 1% | 4% | 4% | 7% |
| **SeqRun07** | raw | 90040 | 91022 | 81026 | 1 | 383072 |
|  | merging | 23% | 25% | 10% | 2% | 99% |
|  | clipping & trimming | 1% | 8% | 0% | 0% | 100% |
|  | filtering | 6% | 1% | 6% | 4% | 10% |
| **SeqRun08** | raw | 52951 | 132500 | 64 | 1 | 993255 |
|  | merging | 14% | 8% | 17% | 1% | 26% |
|  | clipping & trimming | 89% | 24% | 100% | 1% | 100% |
|  | filtering | 49% | 37% | 28% | 0% | 100% |

**Supplemental table 4:**
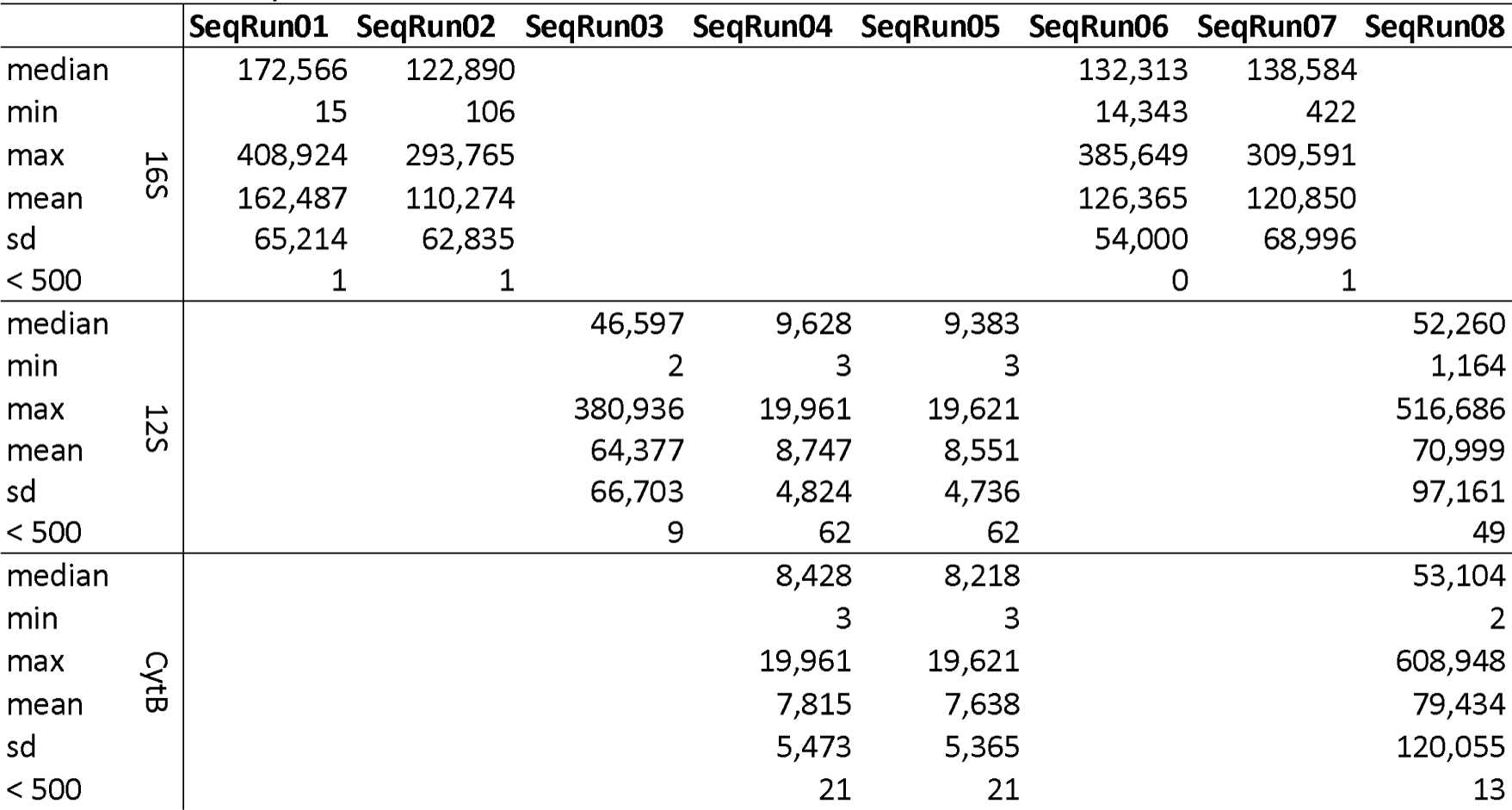
Number of merged R1/R2 reads per sample that were used for the taxonomic assignment for each of the eight sequencing runs. Displayed are the median, minimum, maximum read numbers per PCR replicate, the mean and its standard deviation as well as the number of PCR replicates with less than 500 reads.

